# 2-Locus *Cleave and Rescue* selfish elements harness a recombination rate-dependent generational clock for self limiting gene drive

**DOI:** 10.1101/2020.07.09.196253

**Authors:** Georg Oberhofer, Tobin Ivy, Bruce A Hay

## Abstract

Self-limiting gene drive allows control over the spread and fate of linked traits. *Cleave and Rescue* (*ClvR*) elements create self-sustaining drive and comprise a DNA sequence-modifying enzyme (Cas9-gRNAs, *Cleaver*) that disrupts an essential gene, and a tightly linked, uncleavable version of the essential gene (*Rescue*). *ClvR* spreads by creating conditions in which those without it die because they lack essential gene function. We show that when *ClvR* is implemented in a 2-locus format, with key elements – *Rescue* (and Cargo), and Cas9 and/or gRNAs – located at different genomic positions, spread of the *Rescue* is self-limiting. Drive strength and duration are determined by a recombination rate-dependent generational clock, providing an important point of control for different ecological and regulatory contexts. We implement 2-locus *ClvR* in *Drosophila. Rescue* spreads to high frequency in a Cas9-dependent manner, while the frequency of Cas9 decreases, demonstrating transient drive and loss of future drive potential.

## Introduction

Gene drive occurs when specific alleles are transmitted to viable, fertile progeny at rates greater than those of competing allelic variants. When alleles of genes conferring traits of interest are linked with synthetic genetic elements that mediate self-sustaining drive, spread to high frequency in otherwise wildtype (WT) populations can be achieved for population modification (*1*–*6*) and population suppression (*7*–*9*), forms of population genetic management. These drive mechanisms must be strong enough to spread to high frequency on human timescales, but must also function within diverse and evolving social and regulatory frameworks (reviewed in (*10, 11*)). Central to these considerations are issues of confinement and reversibility: can the spread of transgenes to high frequency be limited to locations in which their presence is sought; can drive be terminated; can new population modifications be exchanged for old ones; and can the population be restored to the pre-transgenic state? Given the diversity of possible scenarios in which drive is being considered, and the competing mandates that any transgene-based technology meant for implementation in the wider world must contend with, no one drive method will be suitable for all occasions, or perhaps even be ideally suited to any particular occasion. Thus, as exploration continues into how gene drive can best be used in the wild it is important to have (in forms that can plausibly be implemented) a diversity of gene drive methods with different characteristics in terms of cost to initiate and maintain a modification at high frequency in a target population, and to confine, terminate, modify or reverse these modifications.

Low threshold self-sustaining gene drive mechanisms include Homing and Y-drive, which are strong drivers at low frequency (*12, 13*), and *Medea* ((*1, 14, 15*)) and *Cleave and Rescue* (*ClvR*) ((*5, 6, 16*)), which are weak at low frequency and strong at higher frequencies. All are predicted to be relatively invasive and may (depending on the details of the system and the ecology) be challenging to confine to a target area due to the fact that continuous low level migration of drive element-bearing individuals into neighboring areas results in these areas being seeded with enough transgene-bearing individuals that spread to high frequency occurs (*13, 17*–*21*).

High threshold self-sustaining gene drive mechanisms include various forms of engineered single- or multi-locus toxin-antidote systems (*3, 22*–*28*), and chromosome rearrangements such as translocations, inversions and compound chromosomes (*4, 29*). These systems require that transgenes make up a much larger fraction of the total wild population before gene drive occurs. Below this frequency transgenes are actively eliminated from the population. High threshold mechanisms are more confinable than low threshold mechanisms by virtue of the fact that the threshold frequency needed for drive in neighboring non-target populations is – depending on the details of the system and migration rate – less likely to be achieved (*21, 30*–*36*). Transgenes can also in principle be eliminated from the population if release of WT results in the frequency of transgenics being driven below the threshold required for drive. These positive points not withstanding, the ability of high threshold drive elements to spread to high frequency in a target area – versus being pushed out of it – while avoiding spread to high frequency in neighboring regions, depends on the details of the drive system, and key aspects of the local ecology, such as migration rates, dispersal distance, and the density and fitness of different genotypes in border regions (reviewed in (*37*)), variables that may often be difficult to quantify.

Given the challenges associated with regulating the behavior and fate of self-sustaining drive elements there is also interest in a second family of approaches, which are our focus here, that include a component of gene drive – which can be regularly reinforced through continued releases – but that are also guaranteed by virtue of the genetics associated with their mechanism of action to lose drive potential (the ability to spread a linked Cargo) at a predictable rate. Non-self sustaining (self-limiting) drive mechanisms are attractive because spread of the desired trait is ultimately always limited in time and space, regardless of the presence or absence of specific physical or ecological barriers or levels of migration. Several such systems have been proposed: Split homing endonuclease genes (a split-HEG) and *Killer-Rescue*. Each works by dividing the gene drive element into two physically separate components, one that is driven into the population and is tightly linked to Cargo genes (endogenous alleles or transgenes that use linkage to hitchhike to high frequency), and another (which is typically genetically unlinked, though see below) that does the driving, but has no ability to enhance its own rate of transmission. There is a progressive decrease in population frequency of the driver element as spread occurs that results from the dispersal of individuals, dilution by WT, and loss by natural selection. The resultant loss of drive activity ultimately limits the spread of Cargo regardless of other variables. The fate of the Cargo is then dependent on the rate at which it is eliminated from the population through natural selection, a process that can potentially be enhanced through the incorporation of unconditional or environmental condition-dependent fitness costs into the Cargo-bearing element.

A simple split HEG locates gRNAs (and any other Cargo genes) at the site of cleavage and homing (thereby disrupting the sequence), while Cas9 is located elsewhere in the genome. In this configuration a homing based increase in gRNA/Cargo copy number only occurs when Cas9 activity and the gRNA/Cargo cassette are present in the same individual. Daisy drive uses a similar strategy, but it is stronger and more invasive in neighboring populations because it includes multiple layers of homing (*33, 38, 39*). Split HEGs have been created in *Drosophila* (*40*–*45*) and mosquitoes (*46, 47*), but not yet shown to spread over multiple generations in otherwise WT populations, often due to the formation of cleavage resistant alleles. Implementations of Daisy drive have not yet been described.

In the *Killer-Rescue* system (*48*) there are two unlinked genes, a zygotic toxin (the *Killer*), which serves as the driver, and a zygotic antidote (the *Rescue*), to which Cargo transgenes are tightly linked (below we refer to this system as *Killer-Rescue*/Cargo). When individuals bearing *Killer* and *Rescue*/Cargo (which are unlinked) are released into a WT population, progeny that inherit the *Rescue*/Cargo with or without the *Killer* survive, while those that inherit the *Killer* but not the *Rescue*/Cargo die. This latter activity serves to bring about an increase in the frequency of the *Rescue*/Cargo-bearing chromosome relative to its WT counterpart. Levels of the *Killer* drop over time whenever it finds itself in non-*Rescue*/Cargo-bearing individuals, and in response to natural selection acting on any associated fitness costs. As the frequency of the *Killer* fades, so too does the drive that maintains the *Rescue*/Cargo construct in the population. An implementation of *Killer-Rescue* has recently been described in *Drosophila*, showing that (with some tinkering, required to identify *Killers* and *Rescues* that worked well together) self-limiting drive can be successful (*49*).

Here we describe and implement a novel mechanism for self-limiting drive, 2-locus *Cleave and Rescue* (2-locus *ClvR*). Drive with 2-locus *ClvR* is much stronger (it drives transgenes with or without fitness costs to a higher frequency and for longer duration per unit introduction percent of transgene-bearing individual) than with *Killer-Rescue*. In addition, drive strength, and duration and extent in time and space for a given introduction percent can be dramatically extended in a measured manner – which is still self-limiting – simply by moving the two components close enough to each other on the same chromosome so that they experience some degree of linkage (segregate from each other during meiosis when placed in cis at rates <50 centiMorgan, cM). Finally, we describe an implementation of 2-locus *ClvR* in *Drosophila* and show that when it is introduced into an otherwise WT population, *Rescue*/Cargo spreads to high frequency while the frequency of the Cas9 driving element decreases. Introduction of WT into this modified population results in no further drive, thereby demonstrating that drive is transient.

## Results and Discussion

### 2-locus *ClvR* configurations with independent segregation

A 1-locus *ClvR* element (*5, 6*)) (also known as Toxin Antidote Recessive Embryo (TARE) in a related proof-of-principle implementation (*16*)), which serves as the starting point for this work, is a self-sustaining gene drive element (Fig. 1A). It consists of a DNA sequence-modifying enzyme such as Cas9/gRNAs that disrupts endogenous versions of an essential gene (located anywhere in the genome) in the germline and in the zygote using Cas9/gRNAs carried over from the mother, and a tightly linked version of the essential gene recoded to be resistant to cleavage and ectopic recombination with the endogenous locus (the *Rescue*) (*5, 6, 16*). 1-locus *ClvR* spreads because Cas9/gRNAs create loss-of-function (LOF) alleles (the drive force) that select against those who fail to inherit *ClvR* in LOF homozygotes. In contrast, those who inherit *ClvR* always survive (in the case of a haplosufficient gene) because they inherit the *Rescue* transgene, which is tightly linked to one or more Cargo genes (*Rescue*/Cargo). Gene drive with 1-locus *ClvR* is self sustaining because tight linkage between Cas9/gRNAs (the driver) and *Rescue/*Cargo (the component being driven) results in both sets of components continuously experiencing the drive benefits of LOF allele creation – an increase in frequency relative to that of the non-*ClvR* chromosome. Drive with 1-locus *ClvR* is frequency-dependent, slow and weak at low frequencies, and rapid and strong at higher frequencies. It lacks a release threshold when fitness costs are absent, but acquires one in their presence. When drive occurs, transgenes spread to genotype or allele fixation depending on the location of *ClvR* and the gene being targeted (*5, 6, 16, 50*). Finally, haploinsufficient or haplolethal genes can also be targeted. Higher thresholds for drive are created, but when drive occurs, it still leads to transgene and/or allele fixation (*5, 6, 16, 50*).

**Fig. 1:**
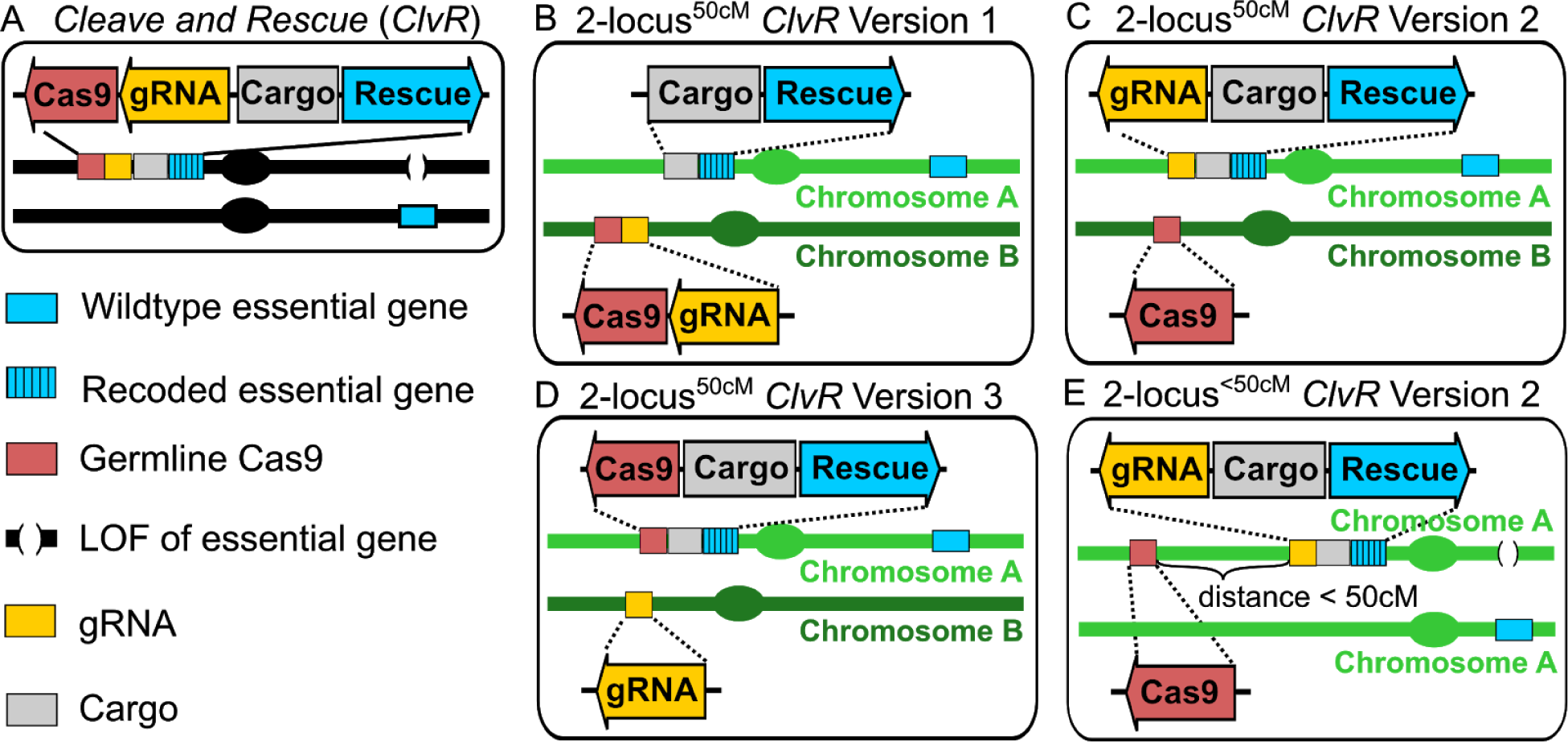
1- and 2-locus *ClvR* configurations. **(A)** 1-locus *ClvR*. **(B)** 2-locus *ClvR*^50cM^ Version 1, in which Cargo/*Rescue* and Cas9/gRNAs are on separate chromosomes, and thus show independent segregation (map distance of 50 cM (centi Morgan)) during meiosis. **(C)** 2-locus^50cM^ *ClvR* Version 2, in which Cas9 is on a different chromosome from gRNAs/Cargo/*Rescue.* **(D)** 2-locus *ClvR*^50cM^ Version 3, in which gRNAs are on a different chromosome from Cas9/Cargo/*Rescue*. **(E)** 2-locus *ClvR*^<50cM^ Version 2, in which Cas9 is located on the same chromosome as gRNA/Cargo/*Rescue* at a distance of less than 50cM. Other versions of 2-locus *ClvR* (Version 1 and Version 3), in which all components are on the same chromosome at a distance of less than 50cM are shown in Fig. S1.

We now consider cases in which the *Rescue*/Cargo and one or both components of germline-expressed Cas9/gRNAs are located on different chromosomes, and thus segregate independently of each other, creating versions of 2-locus *ClvR* designated as 2-locus^50cM^ *ClvR* (cM = centiMorgan). Three configurations are possible. In version 1 (2-locus^50cM^ *ClvR* V1) the Cas9/gRNA construct is located on a different chromosome from that of the Cargo/*Rescue* (Fig. 1B). In versions 2 and 3 (Fig. 1C, D) only one of the Cas9/gRNA components is located on a different chromosome. Because these latter two behave identically, we focus below on 2-locus^50cM^ *ClvR* V2. With 2-locus^50cM^ *ClvR* V1, individuals carrying only the Cas9/gRNA-bearing construct experience essential gene cleavage and LOF allele creation, resulting in death or sterility (and loss of non-*Rescue*/Cargo chromosomes). In contrast, with 2-locus^50cM^ *ClvR* V2 cleavage and LOF allele creation only occurs when *Rescue*/Cargo/gRNA and Cas9 find themselves in the same (viable and fertile) individual. 2-locus^50cM^ *ClvR* V2 (implemented in the experimental section below) is particularly easy to synthesize since organisms carrying each transgene-bearing component are homozygous viable in isolation. In contrast, for 2-locus^50cM^ *ClvR* V1 the Cas9/gRNA transgenes must be introduced into, and maintained within the *Rescue*/Cargo background, as with 1-locus *ClvR* (*5, 6*). Versions of 2-locus *ClvR* in which there is linkage between the components (Fig. 1E, genetic map distance <50cM on the same chromosome) are discussed in a later section.

### 2-locus *ClvR* with independent segregation provides strong, transient drive

We model the behavior of 2-locus *ClvR* using a commonly used framework (*2, 5, 6, 28, 33, 48, 51, 52*) that assumes random mating, non-overlapping generations with no age structure, infinite population size (deterministic populations), and equal and additive fitness costs for each of the transgenes. We use 1-locus *ClvR* and *Killer-Rescue*/Cargo as points of comparison with 2-locus *ClvR* since all utilize a toxin-antidote mechanism of action, and 1-locus *ClvR* (*5, 6, 16*) and *Killer-Rescue*/Cargo (*49*) have been successfully implemented. We begin by showing drive behavior over 300 generations for *Killer-Rescue*/Cargo, and 1- and 2-locus *ClvR*, when introduced at various frequencies, for elements that have no associated fitness costs. When the introduction percent of *Killer-Rescue*/Cargo is low (10-20%), killing of individuals lacking the *Rescue*/Cargo chromosome (which necessarily results in a loss of at least one *Killer* allele), leads to a transient increase in the frequency of the *Rescue*/Cargo-bearing chromosome (Fig. 2A), but the *Killer* is lost before all non-*Rescue*/Cargo chromosomes have been eliminated (Fig. 2B). This results in the frequency of the *Rescue*/Cargo plateauing at some level below allele fixation. At higher introduction percentages, levels of the *Killer* are sufficient to bring about the loss of all non-*Rescue*/Cargo chromosomes. Remaining *Killer* alleles are now protected from loss and float in the population indefinitely, as described previously (*48*). In contrast, with 1-locus *ClvR, Rescue*/Cargo and Cas9/gRNAs spread rapidly to transgene fixation for all introduction percentages because tight linkage of Cas9/gRNAs to *Rescue*/Cargo protects the former from removal in LOF homozygotes and allows it to hitchhike with Cargo/*Rescue* to high frequency, thereby maintaining drive potential indefinitely (Fig. 2C and (*5, 6, 16*).

**Fig. 2:**
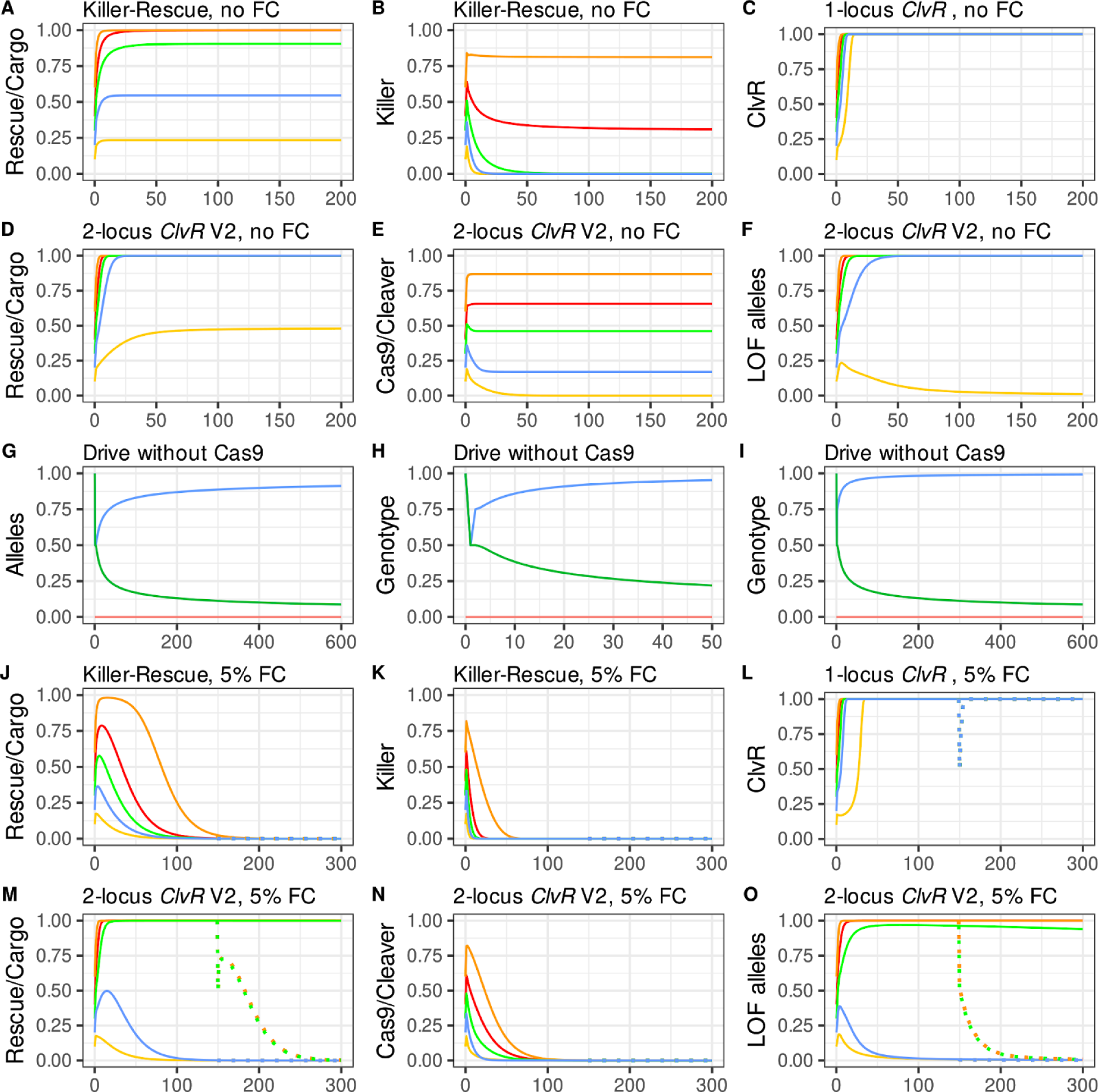
Population dynamics of *Killer-Rescue*/Cargo, 1- and 2-locus^50cM^ *ClvR* V2. **(A-F, and J-O)** Plotted are genotype/allele frequencies (y-axis) over generations (x-axis). Release percentages are 10% (yellow), 20% (blue), 30% (green), 40% (red), and 60% (orange). **(A-F)** elements have no fitness costs; **(J-O)** 5% fitness cost/allele In these panels there is a 50% release percentage of WT in generation 150. Allele and genotype frequencies after this point are indicated with dotted lines. (**G-I)** *Rescue*/Cargo (blue line) allele **(G)** and genotype frequencies over 50 generations **(H)**, and 600 generations **(I);** LOF (green lines); and Cas9 driver (always at zero; red line) in a population initially fixed for Cargo/*Rescue* and LOF alleles, and lacking Cas9 (generation 0), into which a 50% release of WT is carried out in generation 1.

Both versions of 2-locus^50cM^ *ClvR* show an intermediate behavior. Below we present the behavior of Version 2. Fig. S2 shows the behavior of Version 1, which is similar, though not identical. As with 1-locus *ClvR*, drive of the *Rescue*/Cargo chromosome is mediated by the loss of non-*Rescue/*Cargo chromosomes in LOF homozygotes. In the case of 2-locus^50cM^ *ClvR* V1, the driver (Cas9/gRNAs) chromosome brings about the loss of non-*Rescue*/Cargo chromosomes and itself (in sterile or dead individuals) when it creates LOF alleles in the germ cells of those who carry it, and the driver chromosome is not protected by the presence of the *Rescue*/Cargo. This component of 2-locus^50cM^ *ClvR* V1 drive represents the limit of killing and drive by *Killer* in the *Killer-Rescue*/Cargo system. In contrast, with 2-locus^50cM^ *ClvR* V2, LOF allele creation only occurs when the two components are brought together in the same (viable and fertile) individual. Importantly, with both versions of 2-locus^50cM^ *ClvR*, when Cas9/gRNAs or Cas9 are in a *Rescue*/Cargo or *Rescue*/Cargo/gRNA background, respectively, not only are they protected from loss (as with *Killer-Rescue*/Cargo), they still create the LOF alleles at the third, essential gene locus that mediate drive (Fig. 2F). With this “action at a distance”, 2-locus^50cM^ *ClvR* elements create a powerful drive force that can manifest itself in future generations, in individuals that need not carry the driver chromosome (see Fig. S3 for examples). In short, while LOF alleles that mediate drive require Cas9 endonuclease activity for their creation, they exist and segregate independently of it, and do not require its presence (or its loss) for their killing activity. Equally important, the increase in frequency of *Rescue*/Cargo brought about by removal of non-*Rescue*/Cargo chromosomes in LOF homozygotes works to promote the maintenance of LOF alleles in the population because as the frequency of *Rescue*/Cargo increases, the selection pressure that would otherwise bring about the removal of LOF alleles in homozygotes decreases. In consequence, in populations in which *Rescue*/Cargo has been driven to high frequency, LOF alleles can be maintained at high frequency as a latent drive force that manifests *only* when they find themselves in LOF homozygotes that lack the *Rescue*/Cargo chromosome (thereby bringing about further drive of *Rescue*/Cargo), regardless of the current levels of Cas9 and gRNAs.

One way to appreciate the power of this latent drive force is to consider a 2-locus^50cM^ *ClvR* population in which a *Rescue*/Cargo/gRNA with no fitness cost has spread to allele fixation, the Cas9 driver chromosome has been completely eliminated (as would happen during drive if the presence of Cas9 resulted in a fitness cost to carriers, discussed below), and all endogenous copies of the essential gene have been rendered LOF (generation 0 in Fig. 2 G-I). A large number of individuals WT at each of these loci is now introduced into the modified population (release of 50%). Following this introduction the frequency of *Rescue*/Cargo/gRNA and LOF alleles immediately drops. The frequency of LOF alleles continues to decrease over time as natural selection removes them when they find themselves in homozygotes. However, this same force works (transiently) to bring about a substantial increase in the frequency of the *Rescue*/Cargo/gRNA alleles (Fig. 2G) and genotypes (Fig. 2 H, I) since only individuals lacking the *Rescue*/Cargo/gRNA-bearing chromosome are eliminated in the homozygous LOF background, a point returned to below in the discussion of drive experiments described in Fig. 7.

**Fig. 3:**
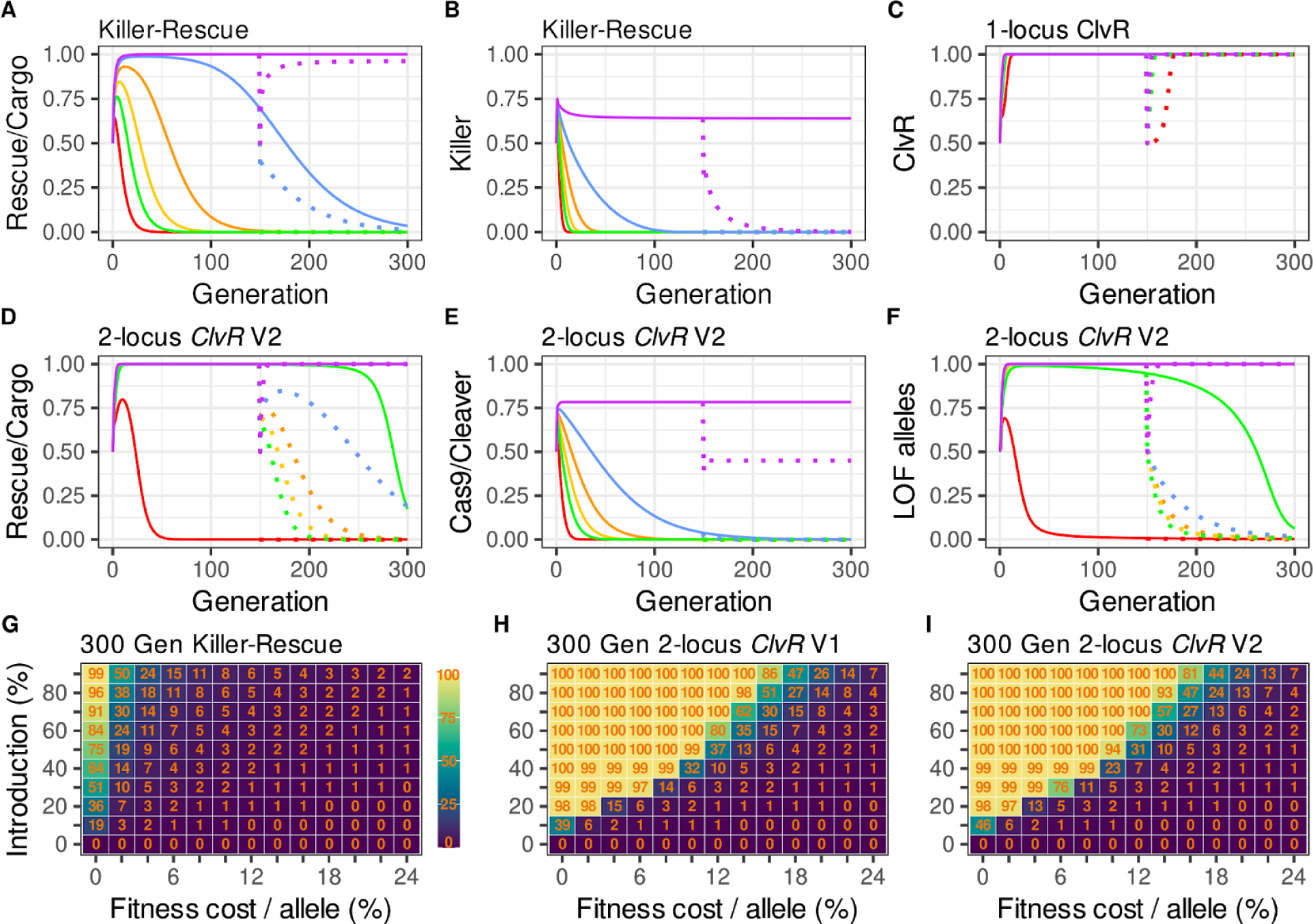
Population dynamics of *Killer-Rescue*/Cargo, 1- and 2-locus^50cM^ *ClvR* for elements with different fitness costs, and introduction percentages. **(A-F)** Population dynamics modeling of different drives introduced at a 50% release percentage, with fitness cost per allele varying from 0-15%. Fitness costs per transgene allele are 0% (purple), 2.5% (blue), 5% (orange), 7.5% (yellow), 10% (green), and 15% (red). Genotype and allele frequencies following initial introductions are indicated with solid lines. Genotype and allele frequencies post WT introductions at generation 150 are indicated in dotted lines. **(G-I)**. Heat maps showing the average *Rescue/Cargo* genotype frequency for the first 300 generations following releases of homozygotes for *Killer-Rescue*/Cargo **(G)**, 2-locus^50cM^ *ClvR* V1 **(H)**, and 2-locus^50cM^ *ClvR* V2 **(I)**, for different introductions and fitness costs/transgene allele. Each rectangle indicates the average Cargo/*Rescue* genotype frequency for the first 300 generations for the introduction and fitness cost associated with the tick marks. Thus, the box in the upper left designates a 90% introduction with a 0% fitness cost.

**Fig. 4.**
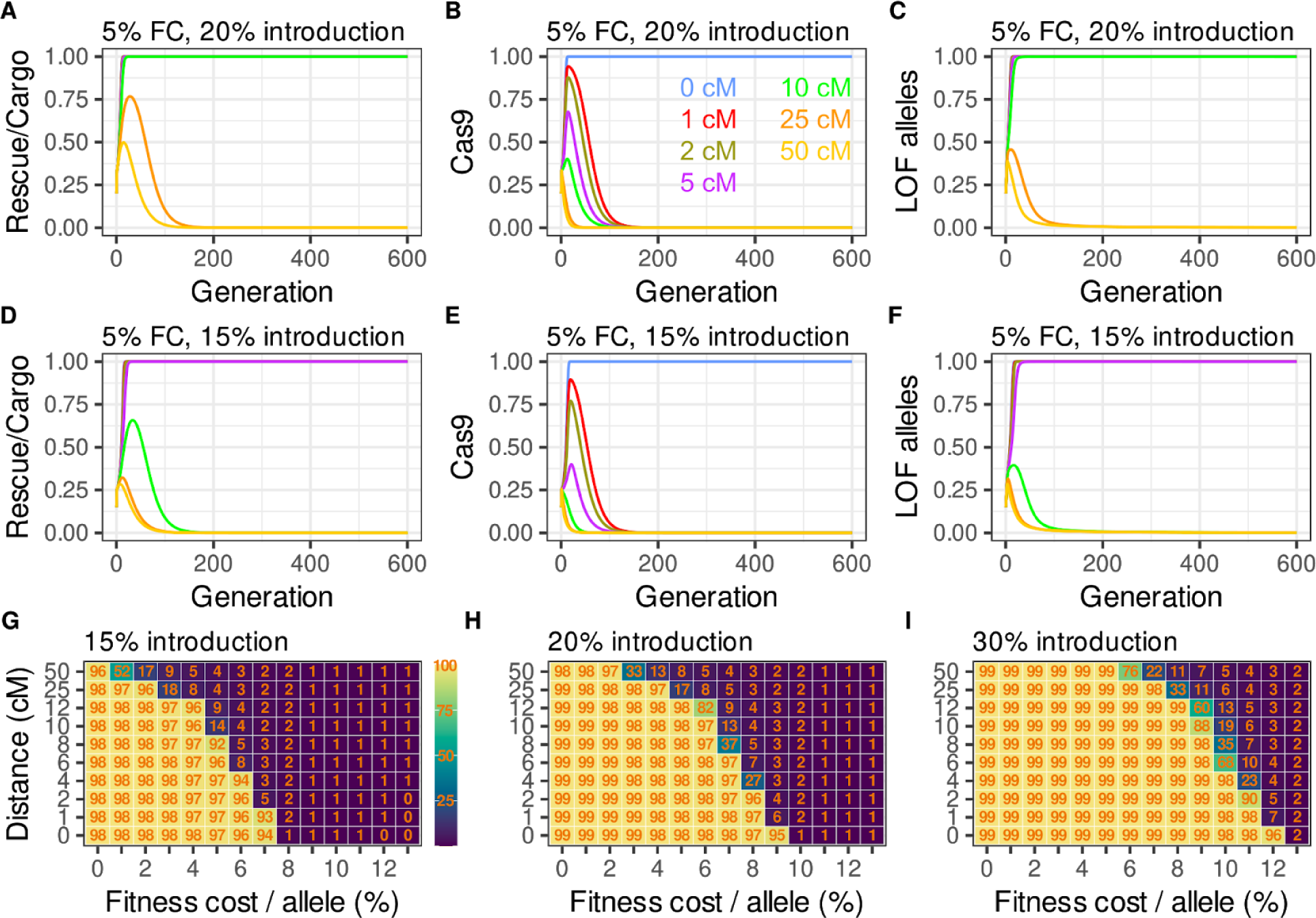
Behavior of 2-locus^<50cM^ *ClvR* V2 with linkage. **(A-C)** Versions of 2-locus^<50cM^ *ClvR* V2, in which the components are separated by different map distances, indicated by colored lines: 50cM (yellow), 25cM (orange), 10cM (green), 5cM (purple), 2cM (brown), 1cM (red) and complete linkage (blue). Each transgene allele has a fitness cost of 5%, and introductions are made at a release percentage of 20%. **(D-F)** As with **A-C**, but with a 15% introduction percentage. **(G-I)** Heat maps showing the average *Rescue*/Cargo/gRNA frequency over the first 300 generations for elements with different fitness costs and map distances between the components (y-axis not to scale), introduced at release percentages of 15%, 20% or 30%.

**Fig. 5:**
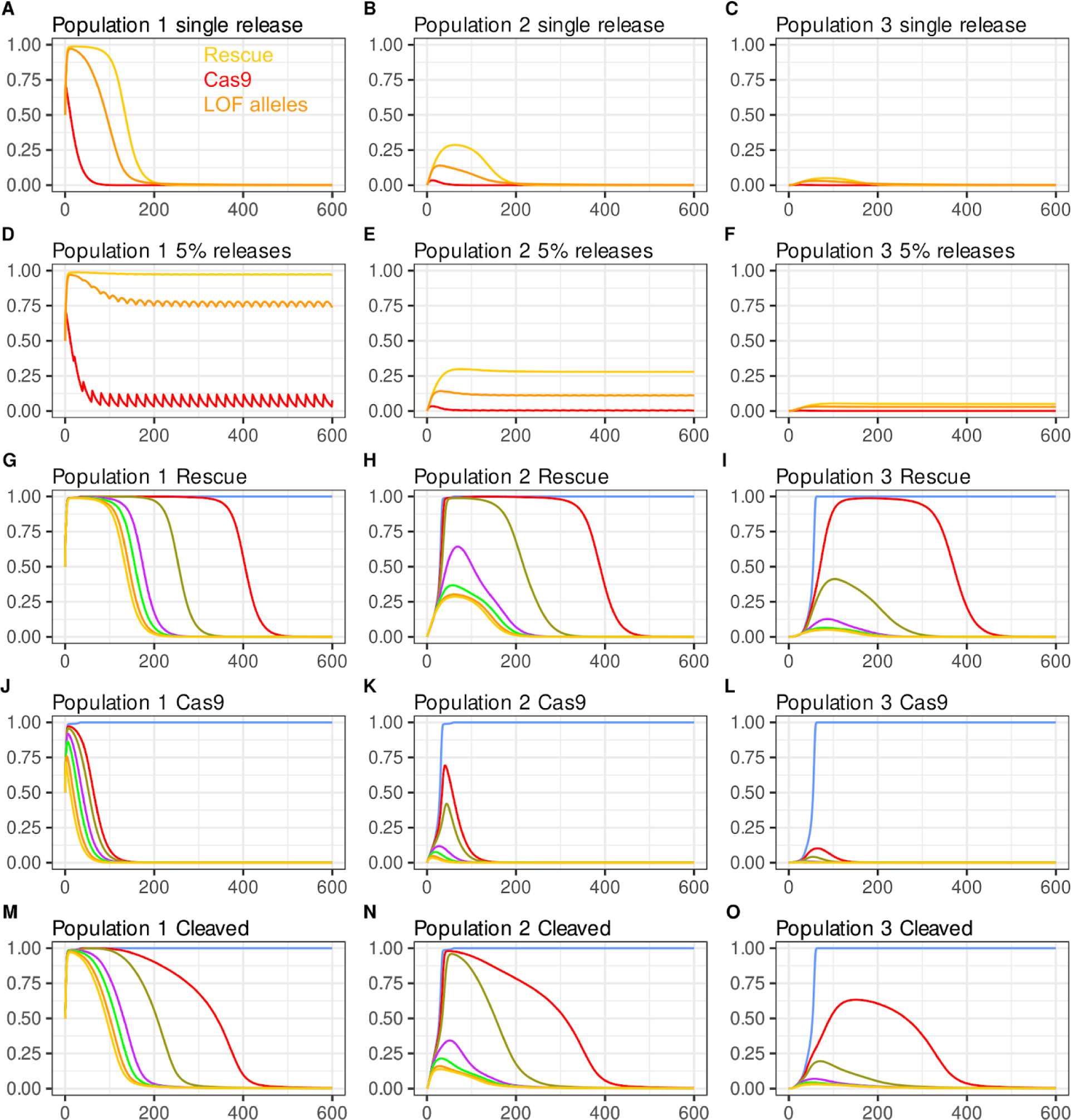
Dynamics of 2-locus *ClvR* in three populations connected by bidirectional migration. Shown are frequencies (y-axis) over generations (x-axis) in 3 populations connected by migration (1% migration rate between populations 1 and 2, and between 2 and 3) after an initial 50% release, for a 2-locus *ClvR* V2 with a 5% FC per allele. **(A-C)** Single release of 2-locus^50cM^ *ClvR* V2. *Rescue*-bearing genotypes (yellow), Cas9-bearing genotypes (red), and LOF alleles (orange). **(D-F)** Same as above but with additional 5% releases every 20 generations. **(G-O)** Single 50% release of 2-locus *ClvR*^≤50cM^ with varying degrees of linkage: 50 cM (yellow), 25 cM (orange), 10 cM (green), 5 cM (purple), 2 cM (olive), 1 cM (red), 0 cM (blue).

**Fig. 6.**
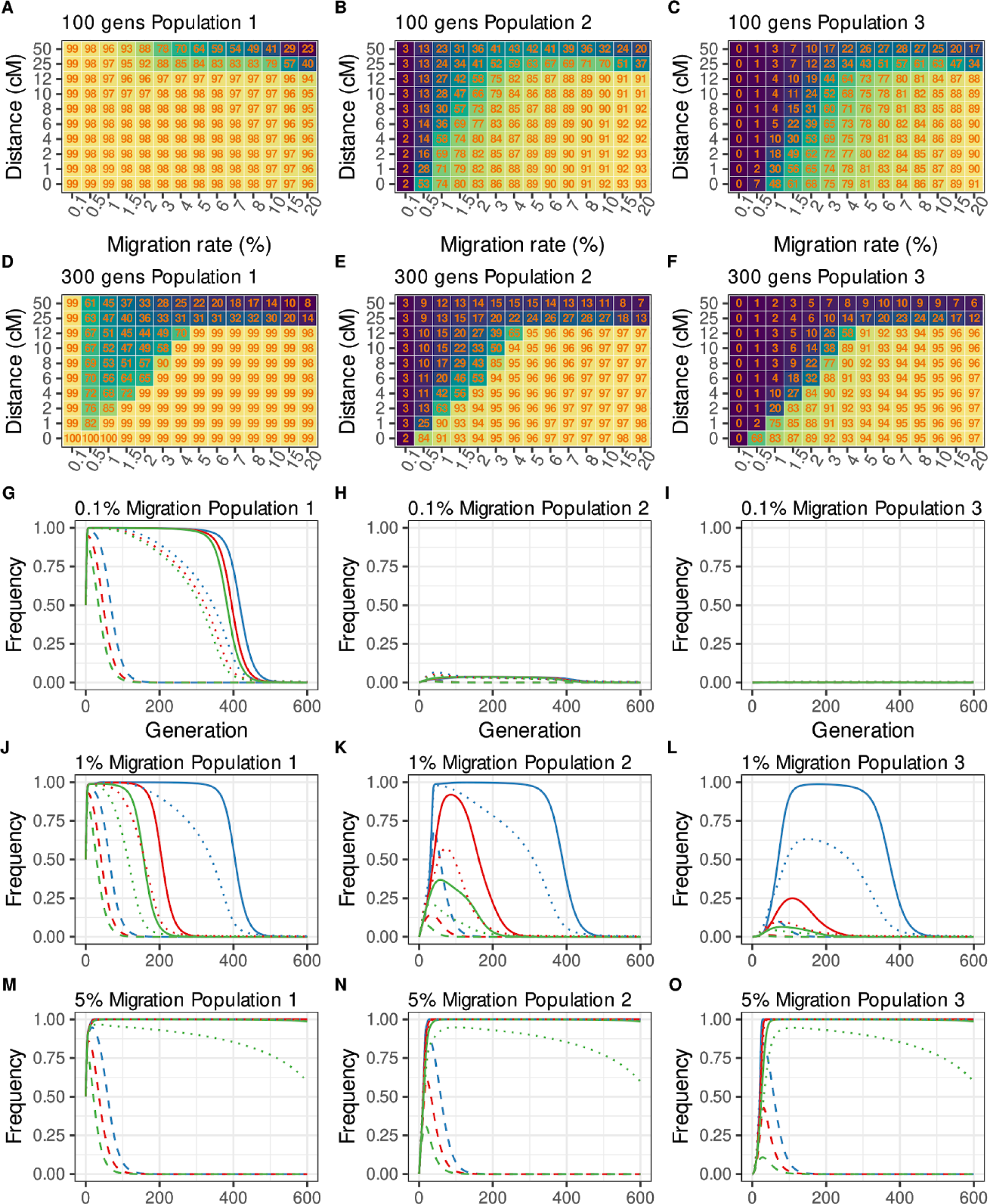
Heatmaps and single modeling runs with different genetic linkage and migration rate. Heatmaps showing the average of *Rescue* frequency for 100 **(A-C)** and 300 generations **(D-F)**. Linkage ranging from 0-50 cM, migration rate from 0.1-20% per generation (plot axes not to scale). **(G-O)** Single modeling runs from above with 0.1% **(G-I)**, 1% **(J-L)**, and 5% **(M-O)** migration rate. 1 cM in blue, 4 cM in red, 10 cM in green, *Rescue* solid, cleaved alleles dotted, and Cas9 with dashed lines

**Fig. 7.**
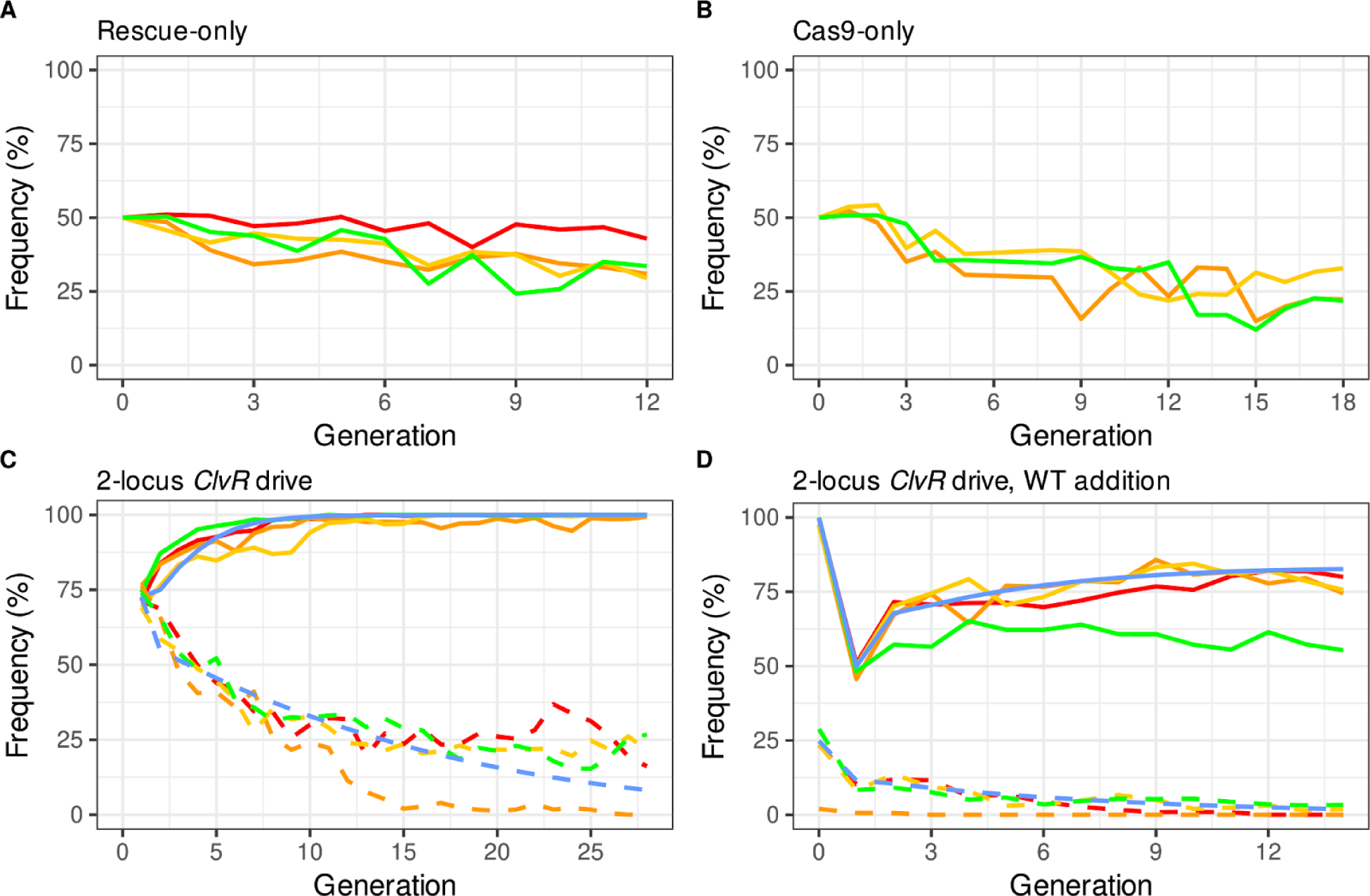
Population behavior of 2-locus drive components *Rescue*^*tko*^ and Cas9/*Cleaver* alone and together as a complete 2-locus^50cM^ *ClvR* V2 element. Behavior of *Rescue*-only **(A)** and Cas9/Cleaver-only **(B)** behavior in a WT (*w*^*1118*^) background. **(C)** Genotype frequencies of *Rescue*^*tko*^ (solid lines) and Cas9/Cleaver (dashed lines) when introduced together as a complete **2-locus**^**50cM**^ ***ClvR* V2 element**, in four replicates (red, green, orange, and yellow). Predicted drive behavior from a model in which *Rescue*^*tko*^ and *Cleaver* have additive fitness costs of 6.5% and 7.5%, respectively, are shown a blue lines (see methods for details). **(D)** Drive populations from **(C)** to which a 50% WT addition was made following generation 15.

Finally, we note that when two independently segregating elements are introduced together into a WT population in double homozygotes (as in these examples), there is transient linkage disequilibrium between the two such that for the first few generations they are found together in individuals more often than would be expected based on their overall population frequencies. Linkage disequilibrium promotes drive of *Rescue*/Cargo with 2-locus^50cM^ *ClvR* because some chromosomes bearing Cas9/gRNAs or Cas9 are transiently protected from death in LOF homozygotes by virtue of an increased frequency of association with *Rescue*/Cargo, while the LOF allele creation that mediates drive continues. In contrast, linkage disequilibrium in the *Killer-Rescue*/Cargo system slows the initial rate of killing and provides no drive benefit, since the killing needed for drive only occurs when both components exist (the *Killer* has not been lost), and the *Killer* and *Rescue*/Cargo have segregated away from each other. As a result of these forces, more non-*Rescue*/Cargo chromosomes are killed per unit of *ClvR* driver chromosome than per unit of *Killer*. This results in higher equilibrium frequencies of *Rescue*/Cargo and the Cas9 driver with 2-locus *ClvR* for any given release percentage of double homozygotes (Fig. 2D,E and Fig. S2 A,B).

We now consider a more realistic scenario in which each transgene-bearing allele results in a 5% fitness cost to carriers, for a total cost of 20% in double homozygotes, for the same range of release percentages (10%, 20%, 30%, 40%, 60%) over 300 generations. In the case of *Killer-Rescue/*Cargo, the *Rescue/*Cargo frequency rises transiently, and then rapidly decays due to natural selection following loss of the *Killer* from the population (Fig. 2J, K). Single locus *ClvR* with comparable fitness costs (10% cost for each combined Cas9/gRNA/Cargo/*Rescue* allele) spreads rapidly to genotype fixation for all introduction percentages (Fig. 2L). Both versions of 2-locus *ClvR* show an intermediate drive behavior that is weak at low introduction percentages, but remarkably strong at higher ones. Thus, when 2-locus^50cM^ *ClvR* V2 is introduced at low frequency (10%, 20%), the frequency of *Rescue/*Cargo/gRNA rises transiently to levels that are higher than those found with *Killer-Rescue*/Cargo (Fig. 2M), and then drops as the frequency of the Cas9 driver drops (Fig. 2N) and the LOF alleles that mediate drive are eliminated from the population through natural selection in homozygotes (Fig. 2O). Version 1 behaves similarly (Fig. S2). However, when either 2-locus *ClvR* is introduced at higher percentages (illustrated for ≥30%), the *Rescue*/Cargo/gRNA spreads rapidly to levels approaching transgene fixation. The frequency of the Cas9 driver decays to zero by ∼ generation 100 (Fig. 2N; Fig. S2E), but *Rescue*/Cargo/gRNA is maintained near transgene fixation for more than 300 generations by the many LOF alleles generated by Cas9/gRNA activity in earlier generations (Fig. 2O; Fig. S2F), which continue to select in LOF homozygotes against individuals that lack the *Rescue*/Cargo chromosome. While this last force is important for maintaining *Rescue*/Cargo/gRNA at high frequency in an isolated population, it is easily subverted through the addition of WT, illustrated for a scenario in which WT are released at a 50% release into the above populations at generation 150 (dotted lines), a point at which the Cas9/gRNA or Cas9 driver chromosome has been completely eliminated. Following an initial drop, *Rescue*/Cargo undergoes a transient increase as non-*Rescue*/Cargo-bearing chromosomes are removed in LOF homozygotes. However, as the frequency of LOF alleles decreases in favor of WT versions of the essential gene this is followed by an inexorable loss of *Rescue*/Cargo/gRNA through natural selection.

To explore 2-locus *ClvR*’s ability to drive in the presence of fitness costs to carriers we consider a scenario in which *Killer-Rescue*/Cargo and 1- and 2-locus *ClvR* V2 are introduced at a constant release percentage of 50%, for a range of element-associated fitness costs (Fig. 3, and Fig. S4 for 2-locus^50cM^ *ClvR* V1). The *Killer* in *Killer-Rescue*/Cargo is able to drive *Rescue*/Cargo to high frequency for ∼80 generations when costs are absent or modest (2.5%/allele), but not when costs are larger (Fig. 3A). These dynamics are reflected in the behavior of the *Killer*, which is rapidly lost except when fitness costs are absent or low (Fig. 3B). 1-locus *ClvR* spreads rapidly for all costs (Fig. 3C). 2-locus^50cM^ *ClvR* V2 spreads to near transgene fixation for >250 generations for all costs except for the highest, 15% per allele (Fig. 3D). The basis for the strong drive by 2-locus^50cM^ *ClvR* V2 can be seen in the extended lifetime of Cas9 as compared with *Killer* (compare Fig. 3B with Fig. 3E, and the concurrent loss of almost all functional endogenous copies of the essential gene (Fig. 3F). The key role LOF alleles play in maintaining *Rescue*/Cargo/gRNA at high frequency can be seen for the scenario involving a 10%/allele fitness cost (green line), in which the decrease in *Rescue*/Cargo/gRNA frequency (Fig. 3D) (long after the Cas9 driver chromosome has been eliminated, Fig. 3E) is preceded by a decrease in the frequency of LOF alleles (Fig. 3F). As also noted above (Fig. 2M), loss of *Rescue*/Cargo/gRNA under conditions that would otherwise support very long-term maintenance at high frequency can be hastened through the addition of WT at generation 150 (Fig. 3D), a point at which the frequency of the driver chromosome has been greatly reduced (2.5% cost/allele) or completely eliminated (costs/allele > 2.5%) (Fig. 3E).

The results presented thus far in Fig. 2 and Fig. 3 identify several conditions under which 2-locus *Rescue*/Cargo/gRNA spends considerable time at high frequency. That said, whenever the presence of *Rescue*/Cargo/gRNA and Cas9 driver chromosomes results in some fitness cost to carriers, and LOF alleles have not spread to allele fixation (see below for the case in which fixation has occurred) due to an earlier loss of the Cas9 driver from the population, LOF alleles and then *Rescue*/Cargo will ultimately be eliminated from the population through natural selection. To get a more general sense of conditions able to support long-term maintenance of *Rescue*/Cargo/gRNA at high frequency we determined the average *Rescue*/Cargo/gRNA-bearing genotype frequency over the first 300 generations for different introduction percentages and fitness costs, an approach used to characterize several other drive mechanisms (*33*). The results of this analysis, for a single release, are shown in Fig. 3G-I, for *Killer-Rescue* and 2-locus^50cM^ V1 and V2, respectively. In brief, *Killer-Rescue*/Cargo is able to maintain *Rescue*/Cargo at high frequency for only a very limited set of conditions that involve a high introduction percentage and very low fitness costs (Fig. 3G). In contrast, both versions of 2-locus *ClvR* drive *Rescue*/Cargo (Fig. 3H) and *Rescue*/Cargo/gRNA (Fig. 3I) to sustained high transgene-bearing frequencies of ≥99% for a large range of introduction percentages and fitness costs.

### Versions of 2-locus *ClvR* that include linkage result in drive with increased strength and duration

Together, the above observations show that 2-locus^50cM^ *ClvR* can provide strong and sustained drive of *Rescue*/Cargo to high frequencies in isolated populations. The introduction percentages required represent a significant fraction of the population. However, these levels are plausible (in at least some cases) as they are substantially lower than those used in earlier nontransgenic insect population suppression programs (*53*). Nonetheless, in contexts where the goal is to modify a population over a large, regional area (an extended area in which multiple target populations are connected by moderate levels of migration, but separated from non-target populations by little or no migration), for a prolonged period, an ideal self-limiting drive system would have lower economic and logistical costs: it would have greater strength at lower introduction percentages, and the drive element would persist and remain active for a longer period of time. In short, drive strength and duration would behave more like that of 1-locus *ClvR*, while remaining self-limited. Here we show how creation of linkage between Cas9 driver and *Rescue*/Cargo components of 2-locus *ClvR* can achieve these goals in a manner in which drive strength and duration are tuned by the frequency of meiotic recombination between the components.

As with 2-locus^50cM^ *ClvR*, there are three possible configurations that involve linkage between 2-locus *ClvR* components. One of these, analogous to 2-locus^50cM^ *ClvR* V2 from Fig. 1C, is shown in Fig. 1E, and the others are shown in Fig. S1. In each configuration (denoted generally as 2-locus^<50cM^ *ClvR*), Cas9 driver and *Rescue*/Cargo components are located in cis, on the same chromosome, and the frequency of meiotic recombination between them is less than 50% (<50 cM). In the case of 2-locus^50cM^ *ClvR*, linkage disequilibrium (the difference between the frequency which alleles of the two components are found together in an individual and that predicted by the product of their allele frequencies), following an initial introduction into a WT population, decays to zero very rapidly. However, when the frequency of recombination between the components is reduced, linkage disequilibrium decays more slowly. During the intervening generations (and as occurs every generation in the context of 1-locus *ClvR*), linkage allows the Cas9 driver element to hitchhike with Cargo/*Rescue* to high frequency. This continues until linkage equilibrium is reached, at which point the Cas9 driver no longer experiences drive. The closer the loci are in cis (on the same chromosome), the more generations it takes for linkage equilibrium to be achieved – thereby creating a generational drive clock. In the limit, drive strength and duration by 2-locus^<50cM^ *ClvR* approaches that of 1-locus *ClvR* as the frequency of recombination approaches zero. However, so long as meiotic recombination between the loci occurs at some rate, linkage equilibrium is always reached, and the duration of drive is limited through the mechanisms discussed above, and below in the context of migration (see discussion of Fig. 5).

These points are illustrated in Fig. 4 A-C, for 7 different 2-locus^<50cM^ *ClvR*s, in which the map distances between the components range from zero (1-locus *ClvR*) to 50cM (2-locus^50cM^ *ClvR* V2); each transgene allele results in a 5% additive fitness cost, and a single release is carried out at a 20% release percentage. Spread of *Rescue*/Cargo/gRNA to high frequency (>99% genotype frequency) fails when the map distance between the components is 50cM or 25cM, but occurs and is maintained for more than 600 generations when the map distances is 10cM or lower. The basis for increased drive with increased linkage can be seen in the plots of Cas9 genotype and LOF frequency. When linkage is present Cas9 undergoes a transient increase in frequency as a result of hitchhiking with *Rescue/*Cargo. The peak frequency reached and persistence time in the population both rise as the frequency of recombination between the Cas9 driver and *Rescue*/Cargo/gRNA decrease. This results in increased LOF allele creation, which creates stronger selection pressure favoring the *Rescue*/Cargo/gRNA-bearing chromosome. However, since recombination between Cas9 and *Rescue*/Cargo/gRNA still occurs, Cas9 and then LOF alleles are ultimately lost through natural selection, bringing an end to drive. A decrease in the introduction percentage to 15% is still compatible with drive to high frequency, but only when drive is stronger – the map distance between the components is 5cM or less (Fig. 4D-F). A general sense of the relationship between map distance and fitness costs, and drive of *Rescue*/Cargo/gRNA to high frequency with 2-locus^<50cM^ *ClvR* V2 elements is shown in Fig. 4G-I, for three different introductions, in which the average *Rescue*/Cargo/gRNA frequency for the first 300 generations is plotted. To summarize, decreasing the frequency of recombination between the components allows lower introductions to be used for elements with equivalent fitness costs, thereby decreasing costs associated with deployment. Decreasing the frequency of recombination also allows elements with higher fitness costs to spread for a given introduction percentage. This feature – the ability to provide extra drive strength for a given introduction percentage (the choice of which will be determined by economics and logistics) – is likely to be important since fitness costs in the wild are probably often underestimated from laboratory experiments. Our modeling shows that when gene drive by 2-locus^50cM^ *ClvR* V2 is considered in a single panmictic population, drive of *Rescue*/Cargo/gRNAs can be achieved for a range of introduction percentages and fitness costs. The strength and duration of drive can be further increased in a controlled manner through the introduction of linkage between the components in the form of 2-locus^<50cM^ *ClvR* V2 elements (See Fig. S5 for behavior of 2-locus^<50cM^ *ClvR* V1 elements under similar conditions). However, so long as recombination between the components occurs drive is always self-limiting.

An important implication of the observations from Figs. 2-4, applied to more realistic, non-deterministic and isolated populations – in which LOF allele fixation can be reached – is that if all WT alleles have been rendered LOF then every member of the population requires the presence of the *Rescue* in *Rescue*/Cargo/gRNA for survival, a state of permanent transgene fixation that is independent of the presence of the Cas9 driver chromosome. The population will remain in this state even when fitness costs are present, and the *Rescue/*Cargo is not at allele fixation. This stands in contrast to the behavior of the *Rescue*/Cargo in a *Killer-Rescue*/Cargo system, in which, if a fitness cost-bearing *Rescue*/Cargo is present at anything less than allele fixation it will be lost through natural selection. Similar considerations apply to the behavior of a Cargo in a split HEG or Daisy drive. In each of these latter systems, in the absence of the driver chromosome there are no forces that work in the same way that ubiquitous LOF alleles do to prevent *Rescue*/Cargo/gRNA loss over time. This ability of LOF allele fixation to hold a 2-locus *ClvR* population at *Rescue*/Cargo transgene fixation (in an isolated population), independent of the presence of a driver chromosome, creates a unique and reversible system by which long-term study of the genetic and ecological impact of population modification can be explored. Following drive, *Rescue*/Cargo transgenes can be held at genotype fixation indefinitely, but they can also easily be eliminated by natural selection (provided their presence results in some fitness cost) following the addition of WT (see Fig. 2 and 3).

### 2-locus *ClvR* and migration

Broadly speaking, there are two general contexts in which population modification with 1- or 2-locus *ClvR* will be carried out. In the first, considered above, the target population encompases the entire range of the species, or multiple populations exist but target and non-target populations are separated by strong barriers to migration, such that the target population can be considered as a single population (because drive is negligible when *ClvR* is present at very low frequency (*5, 6, 16, 50*). In the second, considered below, the target population is linked to non-target populations by significant levels of migration. Here we focus specifically on bidirectional migration. Important questions in this context are how the influx of WT influences the ability of 2-locus *ClvR* to persist at high frequency in the target population, and what the consequences of drive in the target population are for transgene accumulation in non-target populations.

To begin to explore these issues we follow, as an example, the behavior of 2-locus^50cM^ *ClvR* V2 in a 3 population model in which a single introduction into population 1 (the target population) is made at a release of 50%, with a fitness cost per allele of 5% (conditions used in some of the single population scenarios of Fig. 3), and bidirectional migration occurs at a rate of 1% per generation between populations 1 and 2 as well as 2 and 3 (no direct link between 1 and 3). The frequency of *Rescue*/Cargo-bearing genotypes peaks at >99% and has a mean value of >90% for the first 100 generations, after which it rapidly decreases (Fig. 5A). The lifetime of *Rescue*/Cargo/gRNA and LOF alleles at high frequency in population 1 in a 3 population model is much shorter than in an isolated population (Fig. 3D, I) due to the continuous back migration into population 1 of WT alleles at all three loci. At the same time, *Rescue*/Cargo, and to a lesser extent Cas9 driver and LOF alleles, are transferred through migration to population 2. Drive in population 2 due to the creation of new LOF alleles is negligible because the frequency of the Cas9 driver is low. However, when *Rescue*/Cargo/gRNAs and LOF alleles are present at high frequency in population 1, they transiently accumulate in population 2 at significant frequencies, of about 28%, and 12.5%, respectively (Fig. 5B). Levels of *Rescue*/Cargo/gRNAs, Cas9 driver chromosome, and LOF alleles are very low in population 3 (Fig. 5C).

These observations highlight a fundamental challenge for self-limiting drive mechanisms such as *Killer-Rescue*/Cargo, Split HEGs and 2-locus *ClvR*: how to keep the levels of *Rescue*/Cargo high in the target population in the face of continuous incoming migration once the initial input of driver chromosomes has been lost through natural selection and dilution into neighboring populations? Here we consider two solutions. The first is to maintain some frequency of the driver chromosome in the target population through repeated introductions. Fig. 5D-F shows an example in which, following the initial release of 2-locus^50cM^ *ClvR* (as in Fig. 5A), further releases are carried out, every 20 generations, at a release percentage of 5%. These keep the frequency of *Rescue/*Cargo/gRNA-bearing genotypes ∼98% indefinitely. They also lead to stable frequencies of *Rescue*/Cargo/gRNA (26%), Cas9 driver (∼1%) and LOF alleles (12.5%) in population 2, comparable to the peak frequencies observed following a single introduction (Fig. 5A). Levels of *Rescue*/Cargo/gRNAs and LOF alleles remain very low in population 3, while those of the Cas9 driver chromosome are negligible.

A second solution to the problem of how to keep the levels of *Rescue/*Cargo/gRNAs high over time in the face of incoming migration is to incorporate linkage between *Rescue*/Cargo/gRNAs and the Cas9 driver chromosome. Hitchhiking of Cas9 with *Rescue*/Cargo drives Cas9 to higher frequencies in population 1. It also drives an increase in Cas9 frequency in neighboring populations when *Rescue*/Cargo and Cas9 are transferred while still linked in cis (the probability of which is inversely related to the recombination frequency and number of generations since introduction). Increased levels and persistence of Cas9 in target and non-target populations drive the continued creation of LOF alleles, which maintain *Rescue*/Cargo/gRNA at high frequency. These points are illustrated in Fig. 5G-O, for a three population model in which versions of 2-locus^<50cM^ *ClvR* V2 having different recombination frequencies between the components are introduced into a WT population at a release percentage of 50%, with each transgene allele resulting in a 5% fitness cost to carriers. Single locus *ClvR* spreads to transgene fixation in all populations (Fig. 5G-I). Versions that incorporate some degree of linkage have prolonged, albeit still finite, lifetimes at high frequency in population 1 as compared with 2-locus^50cM^ *ClvR* V2 (<100 generations). Linkage also results in increased levels of *Rescue*/Cargo/gRNAs, Cas9, and LOF alleles in populations 2 and 3. This is particularly marked for 2-locus^<50cM^ *ClvR*s in which the recombination rate between the components is 2% or less. Nonetheless, in each case drive is ultimately limited by the movement of WT alleles at the essential locus into the target population, which result in the elimination of LOF alleles (the driver) in the target population through natural selection.

Are there general rules that can be inferred about the relationship between migration rate and recombination distance, and drive dynamics in target and non-target populations? To explore these questions we consider 2-locus *ClvR* V2 elements with a 5% fitness cost per transgene, introduced at a constant percentage (50%) into population 1 of the 3 population model considered above. Bidirectional migration rates vary, as does the recombination frequency between the components. Populations are characterized with respect to average *Rescue*/Cargo/gRNA frequency over the first 100 (Fig. 6A-C) and 300 (Fig. 6D-F) generations, and for allele and genotype frequencies in a representative subset of specific conditions Fig. 6G-O. When the migration rate is very low (0.1% per generation) population 1 behaves as an isolated population, even for 1-locus *ClvR* (Fig. 6A-F). The introduction threshold (due to the presence of element-associated fitness costs) needed for drive in populations 2 and 3 is never surpassed, and thus *Rescue*/Cargo/gRNAs fails to spread in these populations (Fig. 6A-F and G-I). The recombination rate between the components has little effect on the persistence time of *Rescue*/Cargo/gRNAs at high frequency in population 1 (Fig. 6G) since this is ultimately determined by the rate at which LOF alleles are removed in favor of WT (brought in through migration from population 2) through natural selection, long after Cas9 has been eliminated (Fig. 6G-I). In contrast, when the migration rate is somewhat higher (shown here for between 0.5% and 4%) the ability to maintain *Rescue*/Cargo/gRNAs at high frequency in population 1 is strongly dependent on the recombination rate between the components. This is seen most dramatically with the average values for *Rescue*/Cargo/gRNAs frequency at generation 300 (Fig. 6D). When the migration rate is 1%, tight linkage (1cM) is required for long-term maintenance of *Rescue*/Cargo/gRNAs at high frequency. As the degree of linkage decreases (e.g. 4cM and 10cM), so does the average frequency of *Rescue*/Cargo/gRNAs. The basis for this can be seen in Fig. 6J-L. Drive in population 2 begins to wane around generation 100, due to reduced levels of Cas9 and LOF alleles (decreased drive strength) in population 1 and population 2. In consequence, *Rescue*/Cargo/gRNAs never achieves high levels in population 2. This, coupled with the higher rates of back migration, which includes a large number of WT alleles at each locus, results in a more rapid disappearance of LOF alleles. This, in conjunction with element associated fitness costs, drive the frequency of *Rescue*/Cargo/gRNAs down to near zero by generation 300. More generally, the data from Fig. 6A-F show that within the range shown (0.5%-4%) range of migration rates, increased rates of migration must be counterbalanced by decreased recombination frequency (increased drive strength and duration) in order for *Rescue*/Cargo to be maintained at high frequency. This is because incoming WT alleles at the essential gene locus and *Rescue*/Cargo/gRNAs, in the face of waning levels of the Cas9 driver, work to overwhelm drive in population 1.

Interestingly, when migration rates are ≥5%, and the recombination rates are ≤12cM, sustained drive of *Rescue*/Cargo/gRNAs to high frequency occurs in all three populations – they behave as one large population. *Rescue*/Cargo is maintained at high frequency for >300 generations when recombination rates are ≤12cM, and for >600 generations when they are ≤10cM (Fig. 6M-O). Drive to high frequency occurs in all three populations because the rate of movement of Cas9 and LOF alleles into populations 2 and 3 is very rapid, such that drive strength and duration are largely shared between the populations. This can be seen in the similar (though not identical) dynamics of Cas9 and LOF alleles in the three populations, illustrated for specific conditions involving a 5% migration rate, in Fig. 6M-O. Sustained drive to high frequency fails when recombination rates are higher (25cM and 50cM) for the reasons discussed above: the migration of WT alleles at all three loci back into population 1 outweighs the reduced drive associated with rapid attainment of linkage equilibrium, and subsequent loss of of Cas9 and LOF alleles.

The boundary conditions we explored here for long-term persistence are for elements having a specific introduction frequency and fitness cost. Under other conditions these will shift, but the general points about drive behavior are likely to be the same: that the long-term fate of *Rescue*/Cargo in population 1 depends importantly on migration rate. At very low rates the target population behaves as an isolated population. At (and above) some threshold rate, the target and surrounding populations behave roughly as a single population. However, at intermediate migration rates, back migration of WT in the face of dwindling drive in population 1 can dominate, resulting in *Rescue*/Cargo/gRNAs having greatly reduced times at high frequency in population 1. This fate can be offset to some extent by reducing the recombination rate between the components, which increases drive strength and persistence in both population 1 and 2. Thus, while knowledge of the details of local migration rates may not be critical for bringing about initial drive of *Rescue*/Cargo/gRNAs to high frequency, it will be critical to understanding its long-term fate, the scale and timeframe of monitoring needed to ensure adequate ongoing coverage, and the possible need for further releases, either of transgenics to maintain drive, or of WT to limit drive in non-target populations.

These observations provide only a snapshot of 2-locus *ClvR* behavior in space. Nonetheless several general points can be inferred. In the case of 2-locus^50cM^ *ClvR* V2, drive of *Rescue*/Cargo/gRNAs to high frequency in a target population connected to non-target populations by migration can be maintained, but it requires ongoing effort in the form of repeated releases. Alternatively, linkage between *Rescue*/Cargo and Cas9 can be used to increase the time *Rescue*/Cargo spends at high frequency following a single release, particularly when the recombination frequency between the components is low. In either approach *Rescue*/Cargo/gRNAs accumulates to significant frequencies in neighboring populations as a function of the rate of migration (See Fig. 5 and 6). Tight linkage of Cas9 and *Rescue*/Cargo/gRNAs brings about much higher levels of *Rescue*/Cargo in neighboring populations than does repeated release of 2-locus^50cM^ *ClvR* because with tight linkage both loci continue to derive the drive benefits of LOF allele creation: survival (and thus drive) at the expense of non-*Rescue*/Cargo-bearing chromosomes. Finally, the long-term fate of a 2-locus *ClvR* in a target population depends importantly on the details of bidirectional migration.

To summarize, bidirectional migration creates tradeoffs for all 2-locus self-limiting drive systems. *Rescue*/Cargo alleles are free to float in neighboring populations at levels determined by migration rate and fitness, while the corresponding movement of WT into the target population serves to decrease the frequency of transgene-bearing genotypes, particularly when the frequency of the driver chromosome is low. The migration-mediated “passive diffusion” of drive components into neighboring populations contributes to the self-limiting nature of drive, by diluting the Cas9 and LOF alleles needed for drive as spread of *Rescue*/Cargo takes place. However, by its nature this same behavior (which also contributes to the self-limiting behavior of other 2-locus self-limiting drive methods: *Killer-Rescue*/Cargo, Split HEGs) means that drive with these systems does not (unless migration rates are very low, see above) result in sharp borders with respect to gene flow into non-target areas since the Cargo component is free to persist – and in the case of 2-locus *ClvR* with tight linkage, and with Daisy drive (*33, 39*) – continue to drive for some number of generations in non-target populations. Thus, 2-locus drive mechanisms (including 2-locus *ClvR*) are probably best suited to environments in which migration rates in and out of the target region are low (∼ ≤0.1%), or in which a significant frequency of transgenes in regions neighboring the target area is acceptable, but more global spread is not (thus the need for a self-limiting system). These characteristics overlap with those of high-threshold drive mechanisms, but may require tolerance for higher levels of local/regional transgene spillover into non-target areas. Modeling that takes into account features such as density dependence, dispersal distance, and spatial structure are required to more fully understand *ClvR* behavior in specific ecological scenarios.

### Synthesis of 2-locus^50cM^ *ClvR* in *Drosophila*

To synthesize 2-locus^50cM^ *ClvR* V2 in *Drosophila* we used Cas9-mediated mutagenesis (see methods) to inactivate the Cas9 gene in flies carrying a single locus *ClvR* element on the 3rd chromosome (68E) that has the X-linked gene *tko* as its essential gene target (*ClvR*^*tko*^) (*5*). Cas9 mutants in *ClvR*^*tko*^ were created by injecting into heterozygous *ClvR*^*tko*^*/+* embryos a Cas9-RNP-complex preloaded with two gRNAs targeting the Cas9 coding sequence. Heterozygous females carrying an intact *ClvR*^*tko*^ element give rise to >99% *ClvR*-bearing progeny (*5*). Mutants in which Cas9 was mutated to LOF were therefore identified by outcrossing adult female progeny of injected *ClvR*^*tko*^*/+* embryos to WT males (*w*^*1118*^) and looking for Mendelian inheritance of the dominant marker carried within *ClvR*^*tko*^. Several such females were identified, and progeny from one, which also showed Mendelian transmission in outcrosses to WT, were used to generate a homozygous stock used for subsequent experiments, referred to as *Rescue*^*tko*^. The Cas9 open reading frame in these flies includes a 44 bp deletion at the target site of gRNA1, resulting in a premature STOP codon at amino acid 1339. This truncates the PAM-interacting domain (*54*) at the C-terminus of Cas9 by 30 amino acids (Fig. S6E). The *Rescue*^*tko*^ insertion still carries gRNAs targeting endogenous *tko*, the recoded *Rescue*, and a cargo in the form of the dominant marker gene *OpIE-2-tomato*, as determined by sequencing.

The second component of the 2-locus *ClvR* system is located on the second chromosome, at 59D3 (attP docking line from (*55*), and carries a gene encoding Cas9 expressed under the control of germline-specific regulatory elements derived from the *nanos* gene (based on (*56*), modified as described in (*57*)), and a *3xP3-td-tomato* marker. This transgene and stocks that carry it are referred to as *Cleaver* (Fig. S6). Stocks homozygous for both components constitute the final 2-locus^50cM^ *ClvR* V2 and are referred to as *Cleaver;Rescue*^*tko*^ (see Fig. S6 and Version 2 in Fig. 1C with *Cleaver* on the 2nd chromosome and *Rescue*^*tko*^ on the 3rd).

### Genetic *behavior of Cleaver;Rescue*^*tko*^ components alone and in combination

As noted above, loss of Cas9 activity in *ClvR*^*tko*^, which creates *Rescue*^*tko*^, results in *Rescue*^*tko*^ being transmitted from heterozygous females to viable progeny in a Mendelian manner. To further demonstrate that this chromosome lacks drive activity we carried out a multi generation drive experiment. *Rescue*^*tko*^/+ males were mated with WT (*w*^*1118*^) females to bring about a *Rescue*^*tko*^ population allele frequency of 25% in the first generation. Four replicate populations were followed for 12 generations (Fig. 7A). The population frequency of *Rescue*^*tko*^ underwent a consistent, modest decrease over time, similar to that of a control element used in our single locus *ClvR*^*tko*^ drive experiments (*5, 6*), which carried the recoded *Rescue*, and a dominant marker, but not Cas9 or gRNAs. Similar drive experiments were performed with the Cas9-bearing *Cleaver* 2nd chromosome. Here, the population frequency of the transgene-bearing cassette also underwent a decrease over time, as expected for an element whose presence results in a modest fitness cost to carriers (Fig. 7B). Finally, the signature genetic feature of a complete *ClvR* element is that when present in a heterozygous female, all surviving progeny should carry the *Rescue* element if cleavage-dependent LOF allele creation at the target locus is efficient in the female germline and in the zygote (non-carriers die because they lack a functional copy of the essential gene). Evidence that the levels of Cas9 expressed from the second chromosome *Cleaver*, along with gRNAs from the third chromosome *Rescue*^*tko*^ are sufficient to create LOF alleles at high frequency comes from results of experiments in which *Cleaver*/*+*; *Rescue*^*tko*^/+ heterozygote females were outcrossed with WT (*w*^*1118*^) males. As shown in Table S1, all progeny were *Rescue*-bearing (n=3093), for a cleavage and LOF allele creation rate of >99.97%.

### Drive performance of *Cleaver*; *Rescue*^*tko*^, a 2-locus^50cM^ *ClvR* V2

Results of the above modeling and experimental tests of *Cleaver;Rescue*^*tko*^ components predict that introduction of *Cleaver;Rescue*^*tko*^ into a WT population should result in drive of the *Rescue*^*tko*^ construct to high frequency. To test this hypothesis we mimicked an all male release by crossing double homozygous *Cleaver/Cleaver;Rescue*^*tko*^*/Rescue*^*tko*^ males to WT *w*^*1118*^ females. These mated females were then combined with mated WT females, and the mixed population was allowed to lay eggs into a bottle for one day and then removed. Progeny were allowed 13 days to develop to adulthood and mate. Adults were then anesthetized with CO2 and scored for the presence of *Cleaver* and *Rescue*^*tko*^ markers. They were then transferred to a fresh food bottle to repeat the cycle. Four replicate populations were followed. From all the scored genotypes we calculated the total number of flies carrying the *Rescue*-linked Cargo (*Rescue*^*tko*^ and *Cleaver;Rescue*^*tko*^; denoted as Cargo-bearing), and Cas9 (*Cleaver/+* and *Cleaver/Rescue*^*tko*^; denoted as Cas9-bearing). Results are plotted in Fig. 7C. Starting from 69%-72% Cargo-bearing individuals in the first generation all four replicates reached >97% *Rescue*/Cargo-bearing by generation 11. At the same time the frequency of the *Cleaver* chromosome slowly decreased to between 30% and 0%. Three of the four replicates, in which Cas9 levels fell to between 26% and 18%, reached genotype fixation by generation 14. In contrast, in replicate B, in which the frequency of the *Cleaver* dropped almost to zero, the frequency of *Rescue*^*tko*^ remained high, but did not stabilize at genotype fixation. Population dynamics of individual replicates are presented in Fig. S7.

Successful population modification should also be reflected by an increase in *Rescue*^*tko*^ allele frequency and a decrease in that of *Cleaver*. We determined these frequencies at generation 25, after *Rescue*/Cargo genotype frequencies had been at or near transgene fixation for 10 generations. 100 males from each of the above drive populations were individually crossed to WT (*w*^*1118*^) females, and the offspring were scored for the presence of dominant markers (heterozygous males produce 1:1 transgenic:WT offspring; homozygous males produce 100% transgenic offspring). As shown in Table S2, the frequency of the *Rescue*^*tko*^ allele increased dramatically, from between 34.6%-38.2%, to between 86.3% and 87.3% for the 3 populations in which *Cleaver* was still present at some level (allele frequencies between 9.3% and 17.5%), and to 83% for the population in which Cas9 was lost. These results are well explained by a model in which *Rescue*^*tko*^ results in a fitness cost to carriers of 6.5%, and *Cleaver* a fitness cost of 7.5%. Fitness costs were estimated by a non-linear least squares fit of our model to our data, (see Methods). That said, it is important to recognize that the estimated costs reflect behavior of these transgenes in a diversity of different genetic backgrounds, which themselves are present at varying frequencies throughout the drive experiment. Thus, they constitute only rough snapshot of fitness.

### Drive by the 2-locus *ClvR Cleaver*; *Rescue*^*tko*^ is transient

Together the above results demonstrate that a 2-locus^50cM^ *ClvR* V2 can be used to drive a *Rescue*/Cargo/gRNA transgene to high frequency, while the frequency of the Cas9 driver undergoes a contemporaneous decrease. How much remaining drive potential do these modified populations contain? To explore this question we took adults from the 4 drive populations above at generation 15 and combined them with an equal number of *w*^*1118*^ WT flies as the seed for the next generation of bottles. Offspring were then characterized as above for another 14 generations. As expected, addition of *w*^*1118*^ resulted in an immediate drop in the frequency of *Rescue*^*tko*^ and *Cleaver* individuals in the first generation. This is due to the presence of progeny from matings between *w*^*1118*^ individuals. In generation 2 the frequency of *Rescue*^*tko*^-bearing individuals increased. This also is expected and reflects the fact that many *Rescue*^*tko*^-bearing individuals in the seed generation were likely to be homozygous, with progeny from matings between them and *w*^*1118*^ now being heterozygous. The fate of *Rescue*^*tko*^ in the subsequent 12 generations depends on several forces. Fitness costs associated with *Rescue*^*tko*^ will drive its frequency down. At the same time, low levels of *Cleaver*, coupled with an initially high frequency of LOF alleles, will work to support a transient increase in *Rescue*^*tko*^ frequency. A transient rise in *Rescue*^*tko*^ frequency is hinted at in several of the replicates. However, what is not observed is a strong and consistent rise in *Rescue*^*tko*^ frequency from generations 3 onwards, as occurs in the presence of high frequencies of the *Cleaver* (Fig. 6C), demonstrating that drive by 2-locus *ClvR* is transient.

## Conclusion

In conclusion, our results show *ClvR* selfish genetic elements can be created that have different characteristics in terms of cost to initiate and maintain a modification at high frequency in a target population. 1-locus *ClvR* requires the smallest releases (and thus lowest cost) to bring about population modification, but is also self-sustaining and relatively invasive. In contrast, all versions of 2-locus^50cM^ *ClvR*, including those with tight linkage, are self-limiting. In consequence, their spread to high frequency in space is also limited and population transgenesis can (and will ultimately) be reversed through natural selection if releases stop and transgenes carry any fitness cost. 2-locus *ClvR* uses a very simple toolkit of three components to bring about drive: a site-specific DNA sequence modifying enzyme such as Cas9 and gRNAs that guide it to specific targets, sequences sufficient to bring about germline expression (which need not be germline specific) of Cas9, and a recoded, cleavage resistant version of an essential gene able to rescue LOF phenotypes generated by DNA sequence modification of endogenous copies of the essential gene. The key components are orthogonally acting – gRNAs, essential genes, and *Rescues* are each highly specific to the genes being targeted and rescued – and are indefinitely extensible (because any essential gene can serve as a target). Results in *Drosophila* show that 1-locus *ClvR* elements (the components of which make up a 2-locus element, just in different locations) that drive to genotype fixation can easily be generated (*5, 6, 16*), and can also bring about cycles of population modification, in which old content is replaced with new (*6*). These features, in conjunction with our modeling showing that the strength and duration of drive can be tuned through incorporation of genetic linkage between the components, argue that 2-locus *ClvR* genetic elements represent a plausible platform for self-limiting gene drive in diverse species and regulatory regimes. Finally, while we have focused herein on describing how linkage can be used to create measured self-limited drive using the components that make up *ClvR*, we note that creation of linkage between the components that make up a split HEG – the Cas9 driver and the gRNA/Cargo that are being driven – can also be used to increase the strength and duration of self-limiting drive in this system.

2-locus *ClvR* V2, in which Cas9 is located at one position and the gRNAs, Cargo, and *Rescue* at another, provides the most useful format for development, testing and implementation of 2-locus self limiting drive. Drive strength and duration are similar to those observed with other arrangements of components (V1 and V3), and the two strains required for drive can be kept separately as homozygous stocks that only show drive when brought together, providing a point of control. The V2 format also makes it straightforward to screen for components that work well together. The keys to success in building 1- or 2-locus *ClvR* elements that drive are to have a high frequency of cleavage and LOF allele creation in *trans* by Cas9 and gRNAs, and efficient rescue of LOF in *cis* by the recoded *Rescue*. The ability of specific transgenes to bring about these activities can be influenced by nucleosome positioning (*58*), the local chromosome environment, and activity of specific gRNAs. With 2-locus *ClvR* V2 the activity of a number of *Rescue*/gRNAs and integration sites can be tested by crossing transgenics to individuals known from other experiments to have high levels of Cas9 expression in the germline. Transheterozygous females should give rise to all transgene bearing progeny in crosses to WT if there is a high frequency of cleavage and LOF allele creation in the maternal germline and early embryo, and 50% of the progeny should survive if the *Rescue* is efficient. Using a similar strategy, the ability of other DNA sequence modifying enzymes that do or do not use double strand breaks to bring about the creation of LOF alleles can be tested at sites where Cas9 is known to work well, following integration using site-specific recombinases or homologous recombination. Transgenes and genomic locations that support high levels of DNA sequence modifying activity can be similarly identified by analyzing the results of crosses between *Rescue*/Cargo/gRNAs at locations known to support efficient killing and rescue (as in this work, from our observations with 1-locus *ClvR*^*tko*^ (*5, 6, 16*) and transgenics expressing the nucleases to be tested. Finally, when the goal is to create 2-locus *ClvR*s with linkage, components can be tested in *trans*, as above, before bringing them into linkage through recombination.

In order to take advantage of linkage to extend the strength and duration of drive the recombination rates between specific regions of the genome must be known. Rates across particular regions can be determined in several ways, depending on the tools available in the species (*59*–*62*). They can vary significantly depending on genomic position, and are subject to variation by sex and in response to environmental factors (reviewed in (*63*)). Recombination rates can also be altered in the presence of chromosome rearrangements that include the involved regions. In particular, when a chromosomal inversion spans a region of interest (in this case containing the Cas9 driver and the *Rescue*/Cargo/gRNA) and meiotic recombination occurs between the two loci in an inversion heterozygote, the only viable gametes are the parental types, Cas9/*Rescue*/Cargo/gRNA and WT. Rare DNA damage and repair events could place a 2-locus *ClvR* in an inversion, in an otherwise WT chromosomal population, thereby creating a self-sustaining 1-locus *ClvR* element. However, events such as this will necessarily happen in a single individual in the population. Earlier modeling of the fate of rare alleles of *Medea* (*14, 15, 20*), a single locus-self sustaining toxin-antidote drive system with mechanistic and population genetic similarities to *ClvR*, as well as modeling of single locus *ClvR* (*5, 6, 16*) show that drive by one or a few individuals is insignificant, with the fate of such an allele approximating that of a new, neutral mutation, which is usually lost (*64*).

A more likely scenario is that the wild population into which a 2-locus *ClvR* with linkage is to be introduced already carries, at some frequency, an inversion chromosome that spans the region carrying the 2-locus *ClvR* components. This would have the effect of increasing the duration of drive (if the inversion were common) since recombination in inversion heterozygotes would not produce viable recombinant gametes. However, drive would not become self-sustaining, since recombination and segregation of these components would still happen in non-inversion heterozygotes, the frequency of which can be determined prior to release. Moreover, because the *Rescue*/Cargo/gRNA selects for the chromosome it resides in (the non-inversion chromosome), there will be strong selection against the chromosome that carries the inversion, which cannot acquire the *Rescue* transgene through recombination. In a related vein, we note that for any chromosomal drive element (*Medea*, 1- and 2-locus *ClvR, Killer-Rescue*, and some versions of homing in which the Cargo does not move with the HEG), the component being driven into the population will drag nearby chromosomal alleles from the donor genome along with it as it spreads, until recombination brings the drive element and these alleles into linkage equilibrium. Possible population effects of an increased frequency of donor alleles linked to the site of *Rescue*/Cargo insertion for the biology of the target population will need to be considered in any population modification strategy.

## Material and Methods

### Generation of a *Cleaver* (Cas9) stock

This construct was derived from plasmid pnos-Cas9-nos ((*56*), Addgene #66208, a gift from Simon Bullock). We replaced the mini-*white* marker with *3xP3-td-tomato*, flanked the *nos* promoter und 3’UTR with gypsy insulators and added an attB site to facilitate integration into the fly genome. Details of the cloning procedure are described in (*57*). The construct was injected in to a fly strain with an attP landing site on the 2nd Chromosome at 59D3 (Bloomington stock 9722, (*55*)) alongside a helper plasmid as phiC31 integrase source (Rainbow Transgenic Flies). Injected G0 flies were outcrossed to *w*^*1118*^ and progeny was screened for eye-specific expression of *td-tomato* to identify transformants. Transgenic F1 were balanced over *CyO* to get homozygotes.

### Generation of the *Rescue*^*tko*^ stock

The *Rescue* stock was based off of *ClvR*^*tko*^ described previously (*5*). This stock has a complete *ClvR* selfish element, including Cas9, gRNAs, and the recoded *Rescue*. To implement 2-locus *ClvR* we decided to ablate Cas9 function from this stock, so that it would contain gRNAs and *Rescue* only. This was done by designing two gRNAs that target the Cas9 ORF (PAM in uppercase; gRNA1: tgattcatcagtcaattacgGGG, gRNA2: gtactgataaggctgacttgCGG) to create a LOF mutation in Cas9. CRISPR guide design (*65*) was done in the Benchling software suite. The two gRNAs were pre-mixed with Cas9 protein (all from IDT) to form RNP Cas9 complexes. Final concentration in the mixture was: Cas9 protein 500ng/ul, gRNA1 50ng/ul, gRNA2 50ng/ul. This mixture was injected into the offspring of homozygous *ClvR*^*tko*^ males crossed to *w1118* females (Rainbow Transgenic Flies). To screen for potential loss of Cas9 activity in the injected offspring, we outcrossed the now *ClvR*^*tko*^/+ heterozygous G0 females to *w1118* males. If the *ClvR* element and thus Cas9 is still functional we expect the progeny of this cross to be 100% *ClvR*-bearing due to germline and maternal carryover dependent killing of offspring that does not carry *ClvR*. If we see normal Mendelian inheritance where only 50% of offspring carry the *ClvR* marker, Cas9 function must have been lost. We recovered several G0 females that had lost *ClvR* activity. We chose one of them to build up a stock to carry out all the experiments in this study. Flies were balanced over TM3,*Sb* to get homozygotes. We also sequenced over the Cas9 ORF in this stock to map the mutation induced by the injection of the gRNA pair targeting Cas9 itself (Fig. S6). Construct sequence files, gRNA sequences, and alignments for this stock were published previously (*5*).

### Crosses to generate a double homozygous 2-locus *ClvR* stock

Homozygous *Rescue*^*tko*^*/Rescue*^*tko*^ and homozygous *Cleaver/Cleaver* flies were crossed to a double balancer *CyO*;TM3,*Sb*. Offspring with genotypes *CyO*;*Rescue*/TM3 and *Cleaver*/*CyO*;TM3 were crossed to each other to give double balanced *Cleaver/CyO*; *Rescue*/TM3 flies. These were crossed to each other to generate the double homozygous stock.

### Female germline cleavage rates

We crossed double homozygous *Cleaver/Cleaver;Rescue*^*tko*^*/Rescue*^*tko*^ males to *w1118* females to get heterozygous females in the progeny. These heterozygotes were outcrossed to *w1118* males and the progeny scored for the relevant dominant markers (see Table S2 and Dataset S1 for counts).

### Gene drive experiments

To start the gene drive experiment, we crossed a double homozygous *Cleaver/Cleaver;Rescue*^*tko*^*/Rescue*^*tko*^ stock to *w*^*1118*^ females. These mated females were mixed with WT *w*^*1118*^ mated females at a ratio of 2:1 and transferred to a fresh food bottle as the drive seed generation 0 for a starting allele frequency of 33%. These flies were allowed to lay eggs for one day and removed from the bottles. After approximately 12 days the next generation of flies had eclosed. Flies were scored for their genotypes on a CO_2_-pad and transferred to a fresh food bottle to continue the cycle.

After 14 generations flies were transferred to a fresh food bottle to continue the drive. However, instead of discarding them afterwards, we added an equal amount of w1118 and transferred them to another food bottle to seed the drive experiment from Fig. 6B with 50% addition of WT. This drive experiment was performed as the one described above. All drive counts are in Data S1

### Computational Model and data fitting

We wrote discrete-generation, population frequency models of our various *ClvR* drives in Python to predict their behavior under various conditions. We assumed random mating between individuals, equal mating access between all individuals within a population, drive behavior strictly occurs during gametogenesis, fitness costs strictly affect survival up to mating, and carryover occurs strictly from females to eggs. For our predictive modeling we varied fitness cost per allele, recombination distance between loci, and introduction frequency of different genotypes, while we used fixed cleavage and carryover rates of 100%.

For our fitness parameter estimation in Fig. 7 we used the minimize function of the lmfit package in Python to do a least squares fit of our model to our data. We estimated that our *Cleaver* and *Rescue*^*tko*^ alleles have ∼7.5% (7.47% +/- 0.74%, 95% CI) and ∼6.5% (6.56% +/- 0.038, 95% CI) fitness costs, respectively, relative to *w*^*1118*^ after performing a least-squares fit of our 2-locus *ClvR* model (using a data-averaged introduction frequency and assuming cleavage and carryover rates of 100%) to the Cas9-bearing and *Rescue*^*tko*^-bearing frequency data in the drive experiment (Fig. 7).

### Imaging and figures

Images of fluorescent marker expression in whole flies (Fig. S6) were taken on a Leica M165FC with an AmScope MU1000 eyepiece camera and a DSRed filter. Composites were assembled in GIMP and rescaled to reduce file size. No additional image processing was performed. Modeling and drive figures were plotted in R with the “ggplot2” package. Color palettes for the heatmaps were from the “viridis” package.

### Fly crosses and husbandry

Fly husbandry and crosses were performed under standard conditions at 26°C. Rainbow Transgenic Flies (Camarillo, CA) carried out all of the embryonic injections for germline transformation. Containment and handling procedures for 2-locus *ClvR* flies were as described previously (*57*), with G.O and B.A.H. performing all fly handling.

## H2: Supplementary Materials

Fig. S1 to S7

Table S1 to S2

Data S1 containing drive counts, drive counts after WT addition, control drive counts, cross assay to determine cleavage to LOF, and assay to determine allele frequencies in the gene drive experiment.

## General

Stocks obtained from the Bloomington *Drosophila* Stock Center (NIH P40OD018537) were used in this study.

## Author contributions

Conceptualization, G.O., T.I. and B.A.H.; Methodology, G.O., T.I. and B.A.H.; Investigation, G.O., T.I. and B.A.H.; Modeling, T.I.; Writing – Original Draft, G.O. and B.A.H.; Writing – Review & Editing, G.O., T.I. and B.A.H.; Funding Acquisition, G.O. and B.A.H.

## Data and materials availability

All data is available in the main text and the supplementary materials. Fly stocks generated in this study are available on request.

## Supplementary Materials

**Fig. S1.**
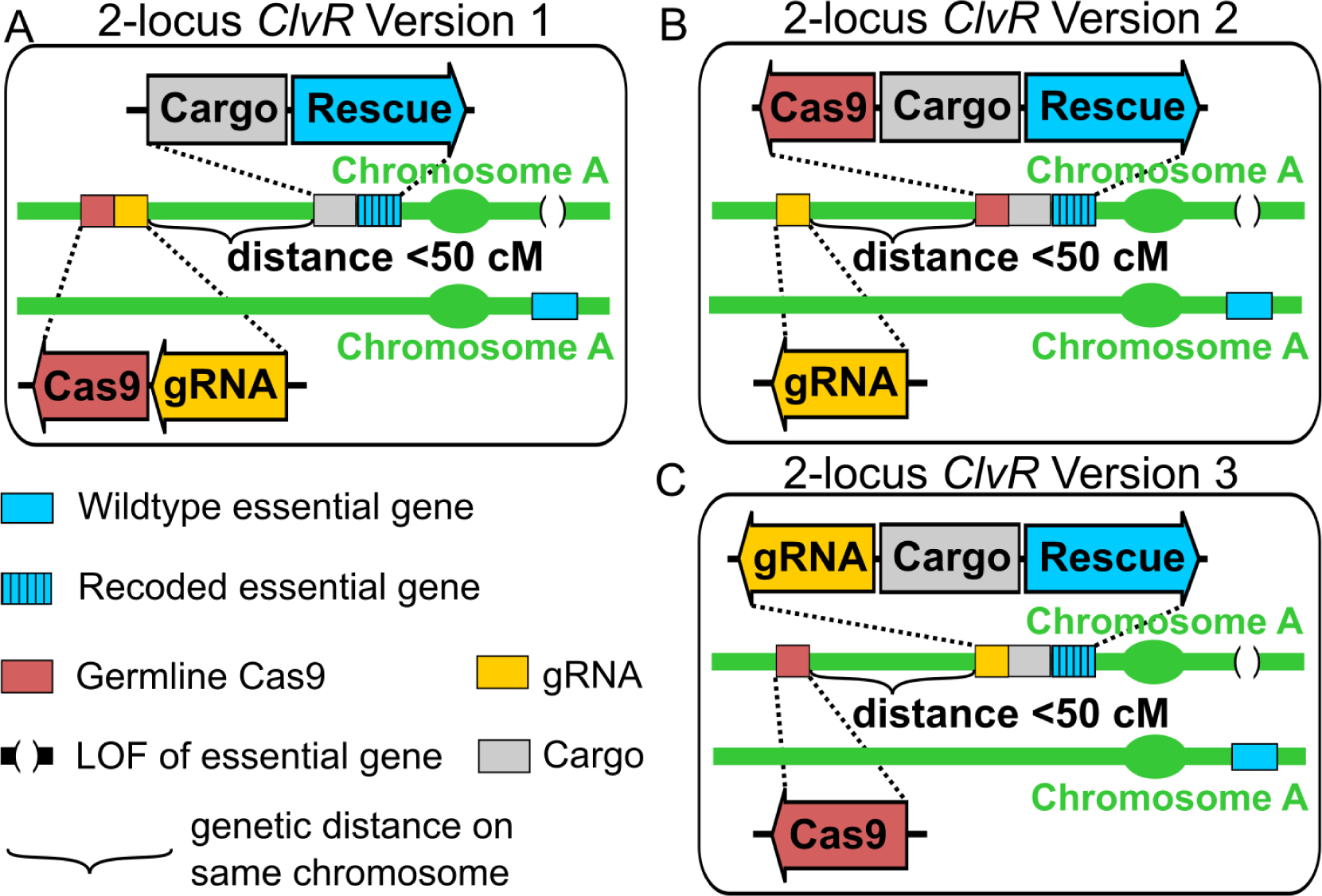
2-locus *ClvR*^<50cM^ configurations. Shown are the possible versions of 2-locus *ClvR* with elements on the same chromosome. See Fig. 1 for 2-locus *ClvR* configurations with elements on different chromosomes.

**Fig. S2.**
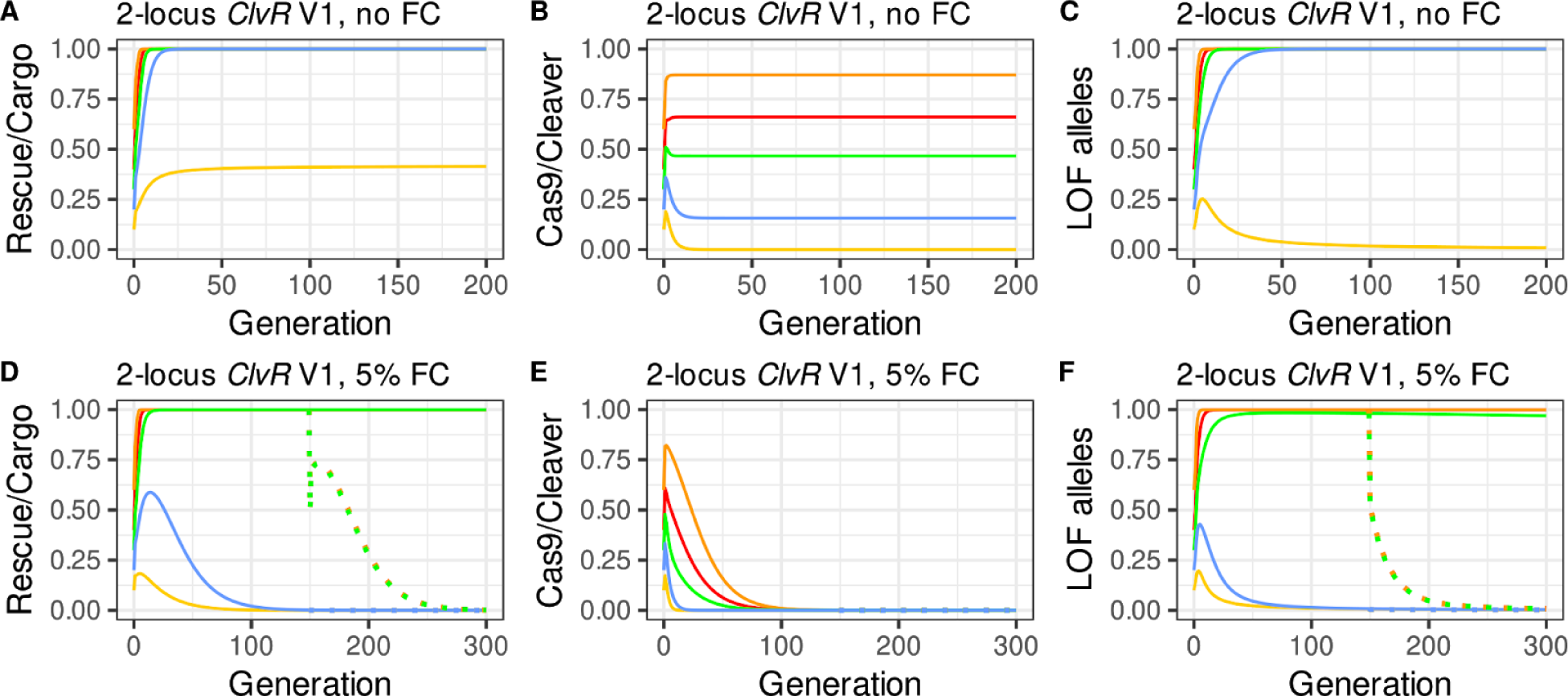
Population dynamics modeling of 2-locus^50cM^ *ClvR* V1 drive introduced at different release percentages. Plotted are genotype/allele frequencies (y-axis) over generations (x-axis). Release percentages are 10% (yellow), 20% (blue), 30% (green), 40% (red), and 60% (orange). In all panels in which fitness costs are present there is a 50% release of WT in generation 150. Allele and genotype frequencies after this point are indicated with dotted lines.

**Fig. S3.**
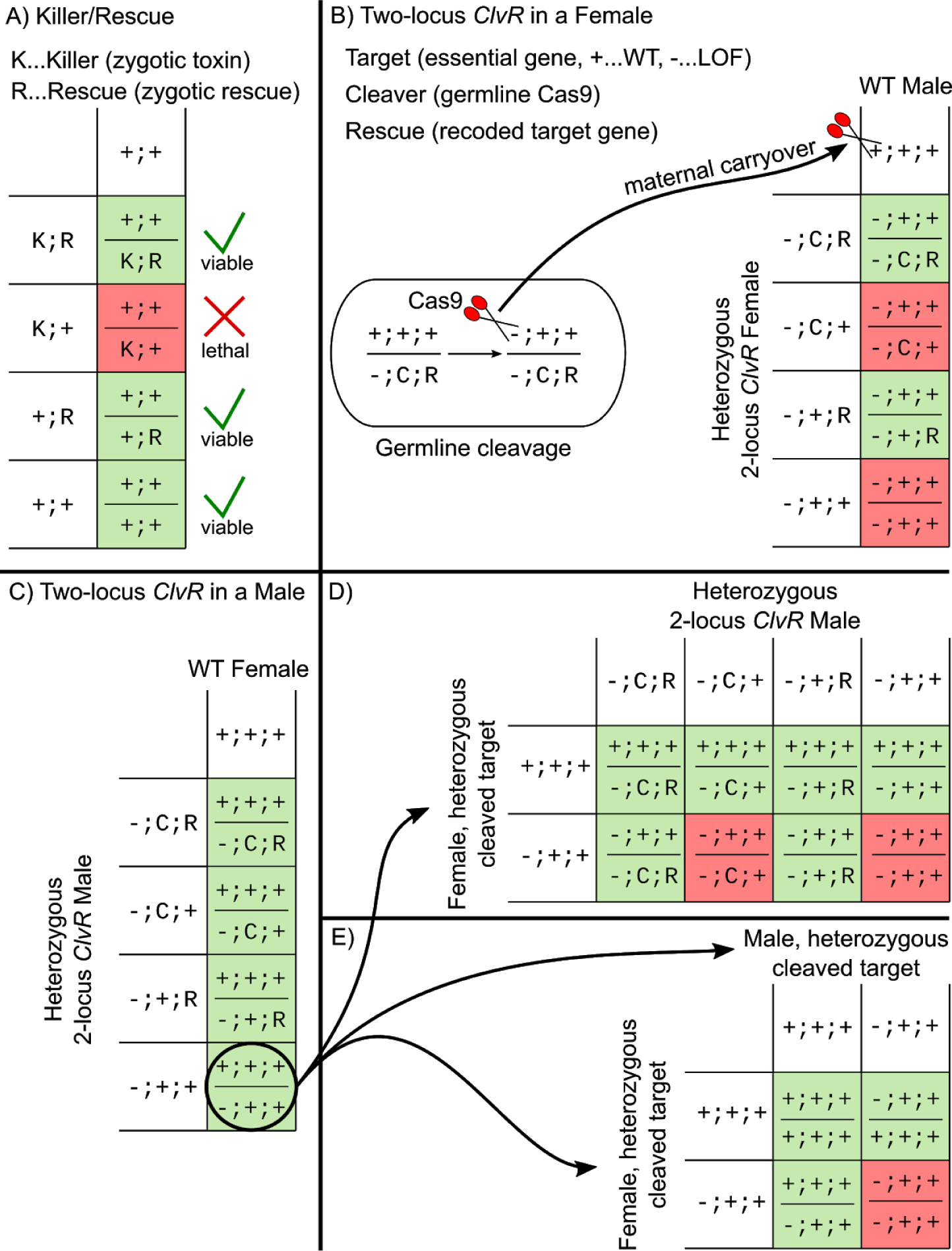
Comparison of *Killer-Rescue* and 2-locus *ClvR* genetics. **(A)** Shown is a cross between a heterozygous carrier of a *Killer-Rescue* to WT. Offspring inheriting only the *Killer* allele die, ⅓ of the remaining offspring carry the *Killer*, ⅔ carry the *Rescue*, ⅓ remains WT. **(B)** Shown is a cross between a female heterozygous for 2-locus *ClvR* to a WT male. Cas9 mutates the target gene to LOF in the female germline. The target allele coming from the WT male gets mutated in the zygote due to maternal carryover of Cas9/gRNA complexes. This results in half of the offspring dying because they don’t carry a copy of the *Rescue.* Of the remaining progeny 100% carry the *Rescue* and 50% carry the *Cleaver*. **(C)** When a 2-locus *ClvR* male mates with a WT female, all the progeny survives. The target gene that was mutated in the male germline remains in the offspring (black circle). **(D)** When an individual heterozygous for the target gene mates again with a *ClvR* male, some of the offspring will end up with 2 mutated copies of the target gene and die. Only individuals that carry the *Rescue* are protected. **(E)** When individuals with one copy of the target gene mate with each other, ¼ of the progeny will die. This results in WT alleles being lost from the population even if the *Cleaver* allele was already eliminated (action at a distance).

**Fig. S4:**
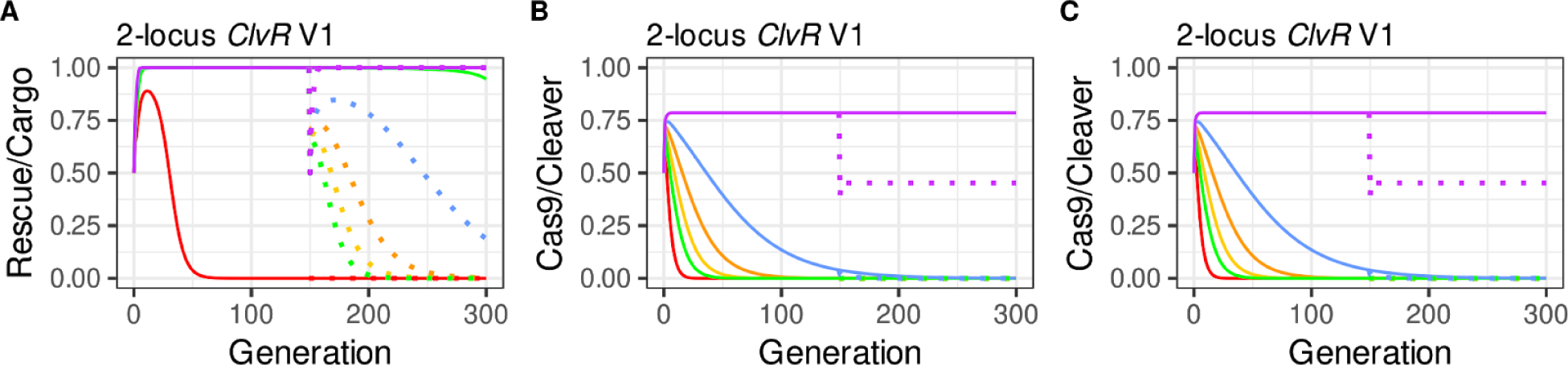
Population dynamics modeling of 2-locus^50cM^ *ClvR* V1 introduced at a 50% release, with fitness cost per allele varying from 0-15%. Fitness costs per transgene allele are 0% (purple), 2.5% (blue), 5% (orange), 7.5% (yellow), 10% (green), and 15% (red).

**Fig. S5.**
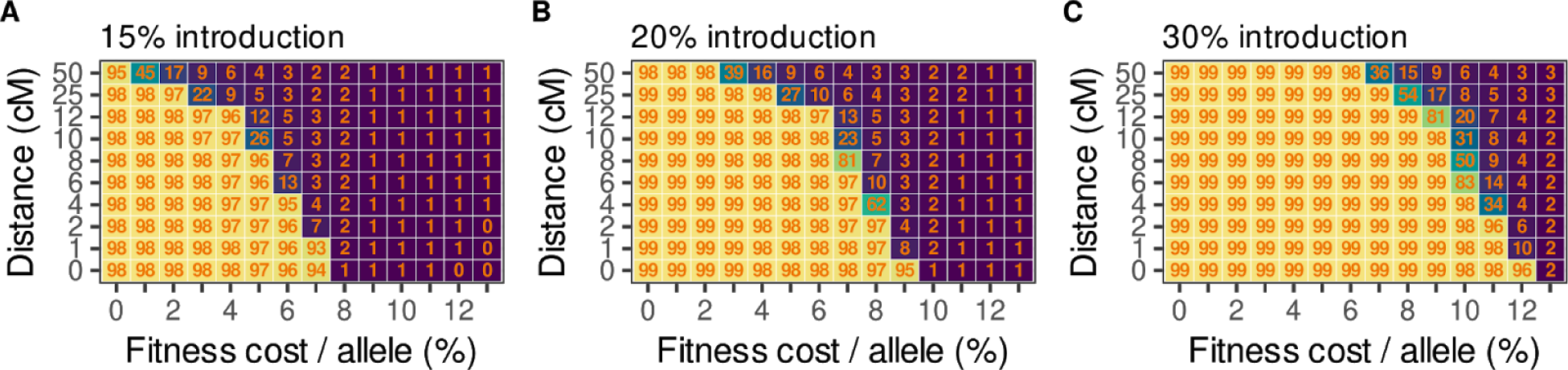
Behavior of 2-locus^<50cM^ *ClvR* V1 with linkage. **(A-C)** Heat maps showing the average *Rescue*/Cargo frequency over the first 300 generations with different fitness costs and map distances between the components (y-axis not to scale), introduced at release percentages of 15%, 20%, and 30%.

**Fig. S6.**
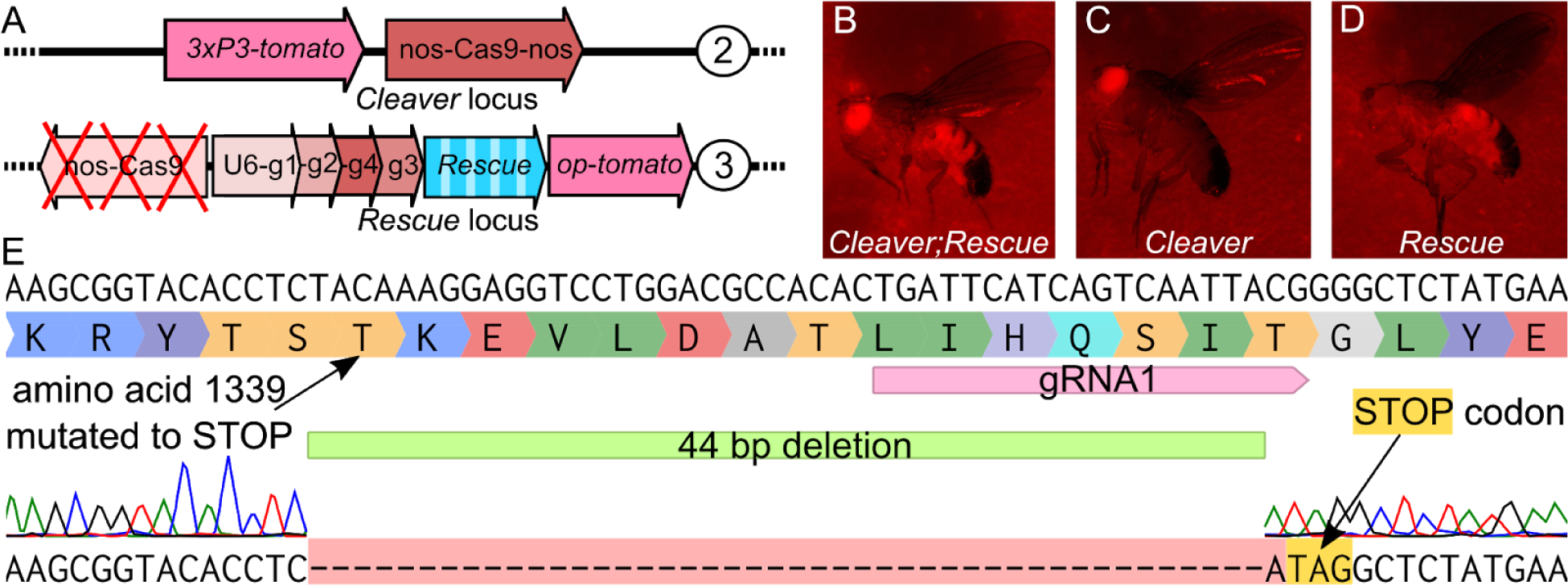
2-locus *ClvR* constructs, markers and alignment of Cas9 mutation. **(A)** Schematic of 2-locus *ClvR* constructs. The Cleaver (Cas9) is on the 2nd chromosome, *Rescue*/Cargo/gRNAs are on the 3rd. **(B-D)** Marker expression in different genotypes. **(B)** *Cleaver;Rescue* fly expressing eye-specific (*3xP3*) and ubiquitous (*OpIE*) *td-tomato*, **(C)** *Cleaver*-only fly expressing eye-specific *td-tomato* **(D)** *Rescue*-only fly expressing ubiquitous *td-tomato*. **(E) Cas9 LOF mutation in original *ClvR***^***tko***^ **locus.** The sequence alignment shows the mutation induced.

**Fig. S7.**
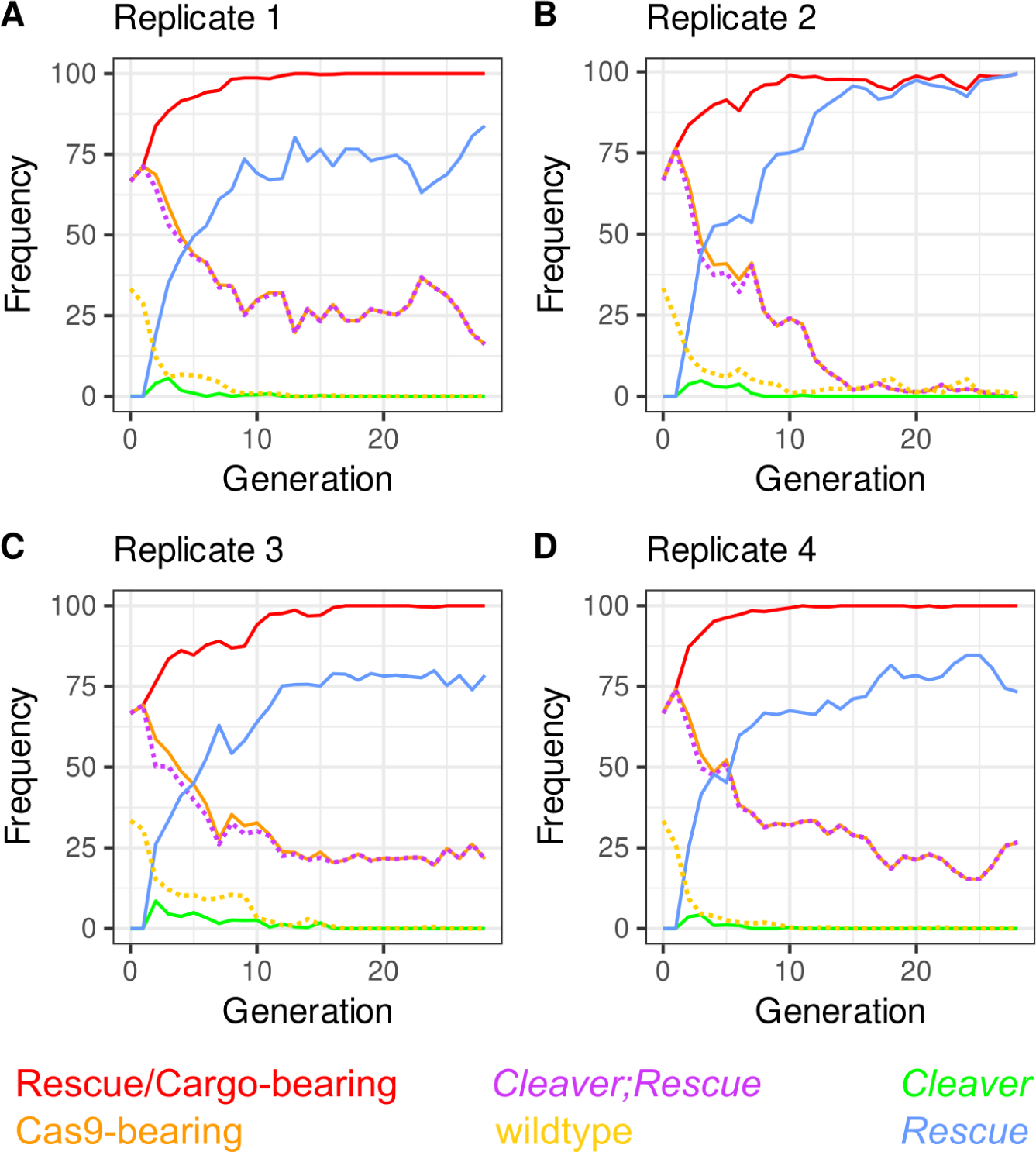
Drive outcomes with all the scored genotypes. Legend on top of panels with *Rescue*/Cargo-bearing in red, Cas9-bearing in orange, *Cleaver/Rescue* in violet, *Cleaver*-only in green, *Rescue*-only in blue, and WT in yellow. **(A-D)** Replicates A-D. WT and *Cleaver;Rescue* in dotted lines for visibility.

**Table S1.**
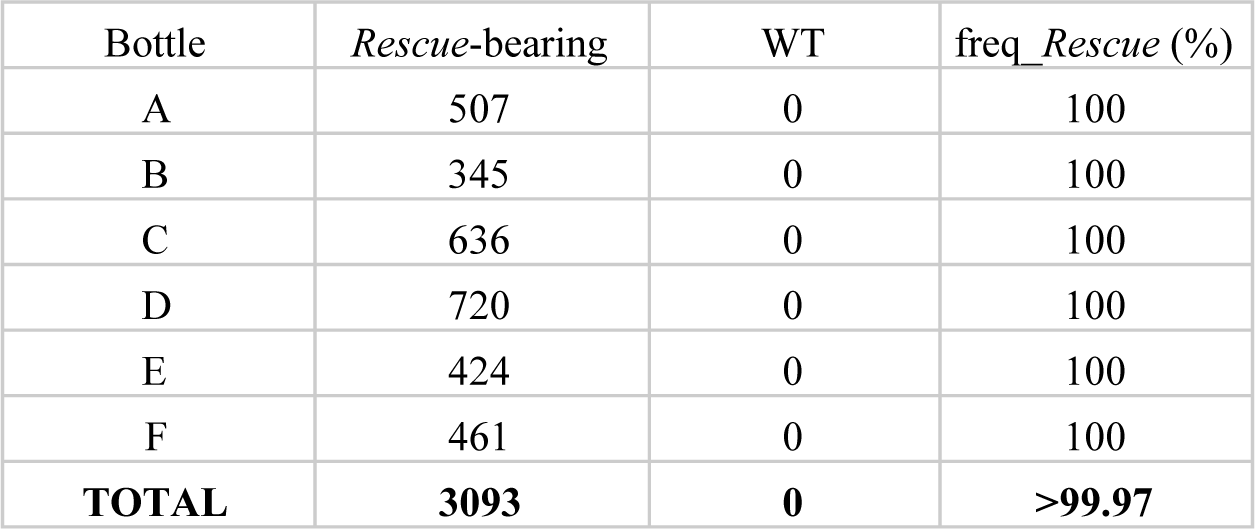
Cleavage rates to LOF in females. Shown are the genotype frequencies in the offspring of a cross between heterozygous *Cleaver/+;Rescue*^*tko*^*/+* virgins and *w*^*1118*^ males. All of the offspring carried the dominant td-tomato marker of *Rescue*^*tko*^.

**Table S2.** Allele frequencies in the drive populations at generation 25. We measured allele frequencies of the drive elements by outcrossing 100 males from the different drive replicates at generation 25 to *w*^*1118*^ virgins and scored the offspring for their respective markers. C= *Cleaver* (Cas9, *3xP3-td-tomato*), R= *Rescue*^*tko*^ (*Rescue*, opie-tomato), +=WT. Examples of how scoring was performed are in Data S1 (allele frequencies).

| Replicate | R/R | R/+ | C/+;R/R | C/C;R/R | C/+;R/+ | total alleles | Cleaver_freq (%) | Rescue_freq (%) |
| --- | --- | --- | --- | --- | --- | --- | --- | --- |
| A | 37 | 19 | 17 | 4 | 3 | 160 | 17.5 | 86.3 |
| B | 58 | 30 | 0 | 0 | 0 | 176 | 0 | 83 |
| C | 43 | 18 | 16 | 0 | 2 | 158 | 11.4 | 87.3 |
| D | 46 | 20 | 14 | 0 | 1 | 162 | 9.3 | 87 |

